# Temporal coding of conceptual distances in the human language area

**DOI:** 10.1101/2025.10.20.683590

**Authors:** Yasuki Noguchi

## Abstract

It is largely unknown how the brain encodes semantic distances between conceptual (verbal) knowledge. I presently show that the information on semantic distances is mainly embedded in a temporal measure (oscillation frequency) of human EEG waveforms. Participants judged a semantic relation between two words (synonym, antonym, thematically-related, taxonomically-related, or unrelated) sequentially presented. Brain rhythms (8-30 Hz) over the left language region, especially the anterior temporal lobe (ATL), were accelerated when a word pair with a high conceptual similarity (e.g. synonym) was presented. Importantly, this change in brain rhythm reflected an inter-word closeness as concepts (holistic similarity, typically seen in synonyms or thematically-related pairs), rather than an inter-word overlap of semantic features (partial similarity, typically seen in antonyms or taxonomically-related pairs). These results highlight a role of temporal measures of neural activity in semantic processing, which explains an inconsistency over previous results using hemodynamic measures.

**Significance statement:** Verbal concepts are connected with each other in the brain, although neural (electric) signals encoding their semantic distances remain unknown. In the present study, I show that the information about inter-conceptual similarity is embedded in speed (not amplitude) of neural oscillatory signals in the healthy human brain. This approach using a temporal measure further revealed the concept-based (not feature-based) processing of word meanings unique to the left language areas.

## Introduction

A fundamental but unsolved issue in neuroscience is where and how semantic knowledge is represented in the brain (Hagoort, 2020; Jamali et al., 2024; Piantadosi et al., 2024). While some researchers proposed a core language network (Binder et al., 2009; Fedorenko et al., 2024) or a semantic ‘hub’ region (Ralph et al., 2017), others reported neural activity of semantic processing distributed across widespread areas outside the core network (Huth et al., 2016; Anderson et al., 2017). Likewise, it is also controversial how the brain handles semantic relationships between words (Popov et al., 2017; Jefferies et al., 2020; Holyoak and Monti, 2021; Li and Pylkkanen, 2021; Wang et al., 2021). Since there are various types of inter-word relationships (Lu et al., 2012; Chiang et al., 2021), they might be processed in separate regions of the brain (Kalenine et al., 2009; Mirman et al., 2017). A dual-hub theory (Schwartz et al., 2011), for example, argues that neural activities in the left anterior temporal lobe (ATL) encode taxonomic relationships between words (connections based on shared categorical features such as animals and foods, e.g. “horse” and “cat”), while those in the left temporo-parietal cortex represent thematic relationships (connections based on co-occurrence in the same scene or event, e.g. “horse” and “saddle”). Although some studies provided the data (partly) supporting this view (de Zubicaray et al., 2013; Lewis et al., 2015; Kumar, 2018; Xu et al., 2018; Thye et al., 2021; Fu et al., 2023; Adezati et al., 2024), others not (Kuchinke et al., 2009; Sachs et al., 2011; Jackson et al., 2015; Chou et al., 2019; Carota et al., 2021; Blackett et al., 2022; Fernandino et al., 2022; Zhang et al., 2023; Liu et al., 2024; Riccardi et al., 2024; Stochel and Sandberg, 2025). How can we resolve those controversies over semantic knowledge and relationships? One way is to develop a new measure for neural activity that has not been used previously. Most studies on the healthy human brain used fMRI (functional magnetic resonance imaging), a measure of which reflected hemodynamic responses. In contrast, neural activity primarily indicates changes in electric signals in a variety of patterns. In case of oscillatory signals like EEG (electroencephalography) and MEG (magnetoencephalography), there are at least two measures to be analyzed; amplitude (power) and speed (oscillation frequency). Changes in hemodynamic (BOLD) signal of fMRI is thought to mainly reflect those in amplitude of oscillatory signals (Murta et al., 2015; Bondi et al., 2025; Xavier et al., 2025). If semantic information is embedded in speed, not amplitude, of neural activity, an approach using a temporal measure would provide a new insight into the problem over neural underpinnings of semantic information.

Based on this idea, here I investigated temporal dynamics of EEG signals underlying semantic processing. In Experiment 1 (**Fig. 1**), neural activities were compared between two representative relations of words; synonym and antonym (Jeon et al., 2009; Scheible et al., 2013; Budel et al., 2023). Previous studies have shown that two words in synonymy (e.g. “increase” and “rise”) are strongly related with each other, as evidenced by a semantic priming effect (Lucas, 2000; Hutchison, 2003) and a high correlation between their word vectors. The strong relationship is also seen in an antonym pair (e.g. “increase” and “decrease”) (Lucas, 2000). It is thus difficult to discriminate synonyms from antonyms solely based on their vector correlations (Samenko et al., 2021). However, while words in synonymy represent virtually the same concept, an antonym pair of words have opposite meanings and thus dissimilar in conceptual space (Budanitsky and Hirst, 2006; Crutch et al., 2012), as evidenced by scores of a similarity rating by human subjects (Hill et al., 2015). For example, “an increase in stock price” and “a decrease in stock price” convey totally different messages that should not be confounded. An antonym is thus regarded as two words that are similar in all but one aspect, in which they are maximally opposed (Willners, 2001; Scheible et al., 2013; Kliegr and Zamazal, 2018). By comparing neural responses to synonyms with those to antonyms, I explored how the information critical to a semantic (conceptual) identity of a word was represented in the brain.

**Figure 1.**
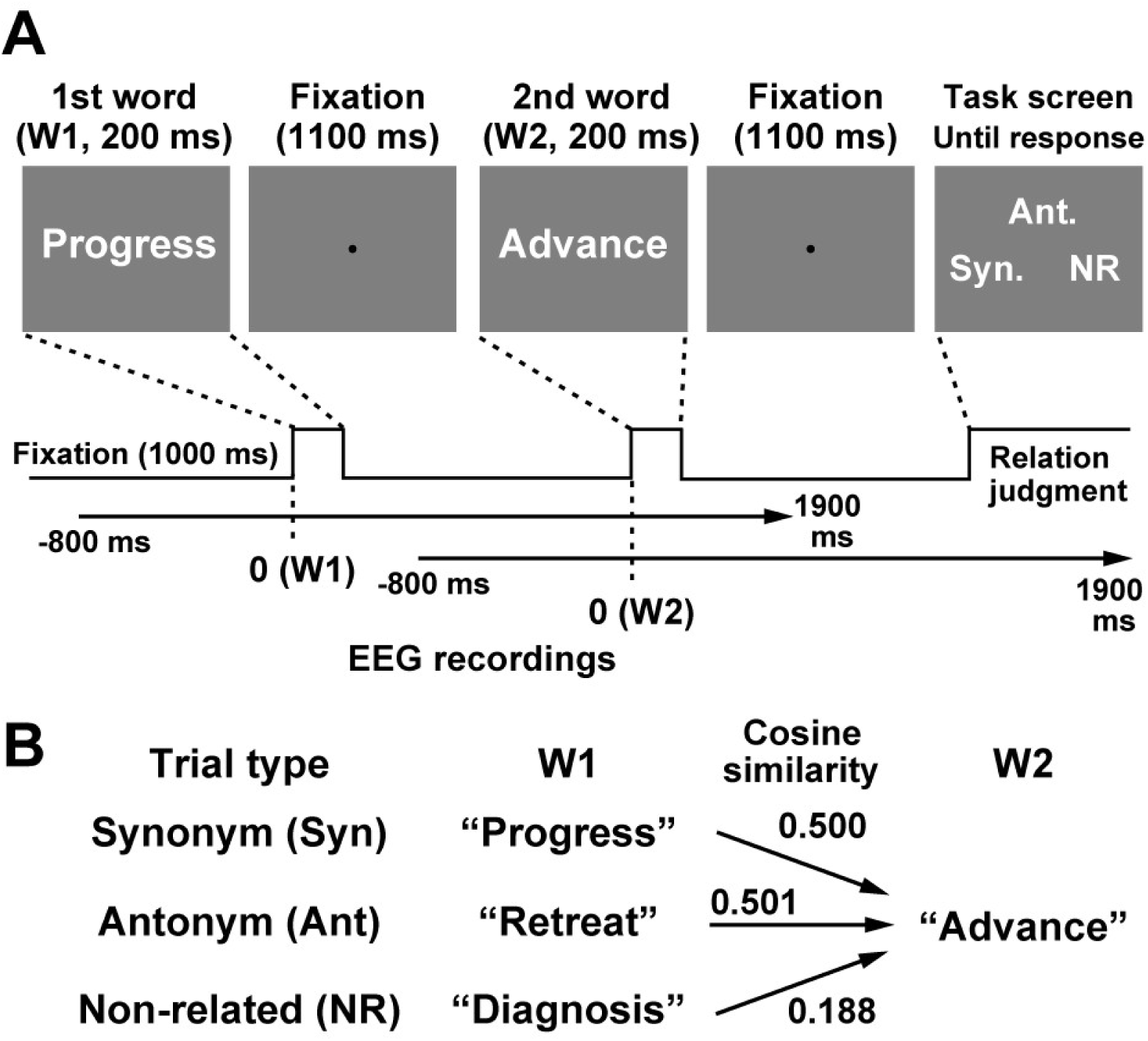
Stimuli and task in Experiment 1. (**A**) Relation judgment (Rel) task. Each trial consisted of a serial presentation of two words (W1 and W2, duration: 200 ms). All words were written in Japanese, although they are translated into English in this figure. Participants judged a semantic relation between W1 and W2 by choosing one of three options (synonym, antonym, or non-related) in the end of each trial (task screen). (**B**) Examples of stimuli in three types of trials. The W1 and W2 in Syn condition had virtually-identical meanings (e.g. “progress” and “advance”), although they had opposite meanings in Ant condition (e.g. “retreat” and “advance”). Trials with non-related pairs of words were also included as a control (NR, e.g. “diagnosis” and “advance”). Those three conditions shared the same set of W2 (e.g. “advance”) so that differential EEG responses to W2 were not attributed to differences in visual inputs over the three conditions.

## Materials and Methods

### Participants

A number of participants (34) was determined with a power analysis using G*Power 3 (Faul et al., 2007). Type I error rate, statistical power, and an effect size were set at 0.05, 0.80, and 0.5 (medium), respectively. In Experiment 1, I measured EEG from 34 healthy participants (native speakers of Japanese, 12 females, age: 18 - 43). Data of four participants were discarded because of a technical issue (missing data, N = 1) and excessive noises (N = 3). Their data were replaced by those of four participants additionally recruited. Laterality quotients (LQs) of the Edinburgh Handedness Inventory (Oldfield, 1971) indicated that 30 participants were right-handed (range: 29.4 to 100) while four participants were not (−100 to −5.9). The same number of subjects (11 females, age: 18 - 36) participated in Experiment 2 (data of two subjects were discarded and thus replaced by additional two). Thirty-two participants were right-handed (LQ: 40 to 100) while two participants were not (−100 and −12.5). All participants had normal or corrected-to-normal visual acuity. An informed consent was obtained from all participants. All experiments were carried out following guidelines approved by the ethics committee of Kobe University, Kobe, Japan.

### Experimental procedures (Exp. 1)

Visual stimuli were generated with the Psychophysics Toolbox for Matlab (Brainard, 1997; Pelli, 1997) and presented on a CRT monitor (refresh rate: 60 Hz). Every participant performed two tasks in separate experimental sessions. In the first (main) task, he/she viewed two words serially presented and judged their semantic relationship (Rel task). A trial started with a fixation point (black circle with a diameter of 0.125 deg) on a gray background for 1000 ms (**Fig. 1A**). This was followed by a sequence of first word (W1, 200 ms), inter-stimulus interval (fixation, 1100 ms), and second word (W2, 200 ms). Both W1 and W2 consisted of two kanji letters (ideograms used in Japan, originally imported from China) vertically arranged.

There were three types of trials; Syn, Ant, and NR (**Fig. 1B**). A synonym pair of words were presented as W1 and W2 in Syn trials (e.g. “progress” and “advance”), while W1 and W2 had opposite meanings in Ant trials (e.g. “retreat” and “advance”). Trials with non-related pairs of words (e.g. “diagnosis” and “advance”) were also included as a control condition (NR trials). Those three types of trials shared the same set of W2 (“advance” in the case above). Across-condition differences in EEG responses to W2 were therefore not explained by differences in visual inputs. In the end of each trial, participants judged a semantic relation between W1 and W2 (task screen). Using a right hand, they pressed a left-arrow key (←), up-arrow key (↑), or right-arrow key (→) in response to the Syn, Ant, or NR trial, respectively. No time limitation was imposed. An experimental session comprised 84 trials in which Syn, Ant, and NR trials (28 trials for each) were intermixed in a random order. Subjects underwent three sessions (252 trials). Correspondences between three response keys (left-, up-, and right-arrow) and three options (synonym, antonym, and non-related) were counterbalanced across the three sessions.

In the second task (Sem task), all 336 words used in the Rel task was individually presented for each trial. Participants answered the meaning of the presented word (target) by choosing one of two options. For example, a target word “Progress” was presented for 200 ms, followed by a fixation screen (1100 ms). Participants then viewed a task screen in which two options were shown on its left and right sides. One option described a correct meaning of the target (“the process of getting better at doing something”), while the other not (e.g. “a sudden event or accident which causes great damage or suffering”). Both options were written in kana (phonograms used in Japan) and kanji letters vertically arranged. A trial was considered as correct when participants chose a correct option by pressing a left-arrow or right-arrow key. Each participant performed three sessions (112 trials for each). An order of the Rel and Sem tasks were counterbalanced across participants.

### Linguistic stimuli

All words in the Rel and Sem tasks were taken from a list of 336 Japanese nouns prepared for the present study. In the Rel task, 252 words were used as W1 (84 for each of Syn, Ant, and NR conditions), while the remaining 84 words were used as W2 shared by the three conditions (**Fig. 1B**). All 336 words were presented in separate trials of the Sem task.

A strength in relatedness between W1 and W2 in the Rel task was strictly controlled by measuring a cosine similarity of word vectors, because those strengths are known to affect neural responses (Teige et al., 2018). I obtained word vectors (size: 1 × 300) for the 336 words from the language database online (fastText library https://fasttext.cc/). Cosine similarities between W1 and W2 (mean ± SD across 84 trials) were 0.500 ± 0.114 (Syn), 0.501 ± 0.126 (Ant), and 0.188 ± 0.072 (NR). A one-way ANOVA over the three conditions indicated a significant main effect (*F*(2,249) = 239.99, *p* < 0.001, *η*^2^ = 0.658). Post-hoc comparisons with the Bonferroni correction showed significant differences in Syn vs. NR (corrected *p* < 0.001) and Ant vs. NR (corrected *p* < 0.001), but not in Syn vs. Ant (uncorrected *p* = 0.95).

It should be noted that word pairs in Syn, Ant, NR trials were balanced in other (visual and phonological) aspects. First, all 336 words in Experiment 1 consisted of two kanji letters, and there was no overlap of letters between W1 and W2. These procedures reduced variations in visual factors over the three conditions, such as numbers of letters and their repetitions. Second, phonological factors were also controlled by equating numbers of moras (phonological units in Japanese) across conditions. Numbers of moras averaged across 84 words (mean ± SD) were 3.63 ± 0.58 (W1 in Syn), 3.55 ± 0.63 (W1 in Ant), 3.57 ± 0.59 (W1 in NR) and 3.57 ± 0.59 (W2). No main effect was observed (*F*(3,332) = 0.3, *p* = 0.83, *η*^2^ = 0.003).

### Analysis of behavioral data

Behavioral data in the Rel task were analyzed along a framework of the signal detection theory. In case of the Syn condition, a trial was considered as hit when participants correctly judged this trial as Syn. A false-alarm (FA) rate was defined as a percentage that participants erroneously judged the other types of trials (Ant and NR) as Syn. A measure of sensitivity (*d’*) was then computed (Snodgrass and Corwin, 1988) using the equation below

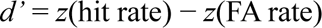

where *z* denotes the inverse cumulative normal function. The *d’*s for the Ant and NR conditions were obtained in the same way. Finally, I conducted a repeated-measures one-way ANOVA with the Greenhouse-Geisser adjustment on the *d’*s over the three conditions.

### EEG measurements

EEG signals were recorded by the ActiveTwo system by Biosemi (Amsterdam, Netherlands), with a sampling rate of 2,048 Hz and an analog low-pass filter of < 417 Hz. I placed electrodes at 32 points over the scalp; FP1, FP2, AF3, AF4, F7, F3, Fz, F4, F8, FC5, FC1, FC2, FC6, T3, C3, Cz, C4, T4, CP5, CP1, CP2, CP6, T5, P3, Pz, P4, T6, PO3, PO4, O1, Oz, and O2 (**Fig. 2A**).

**Figure 2.**
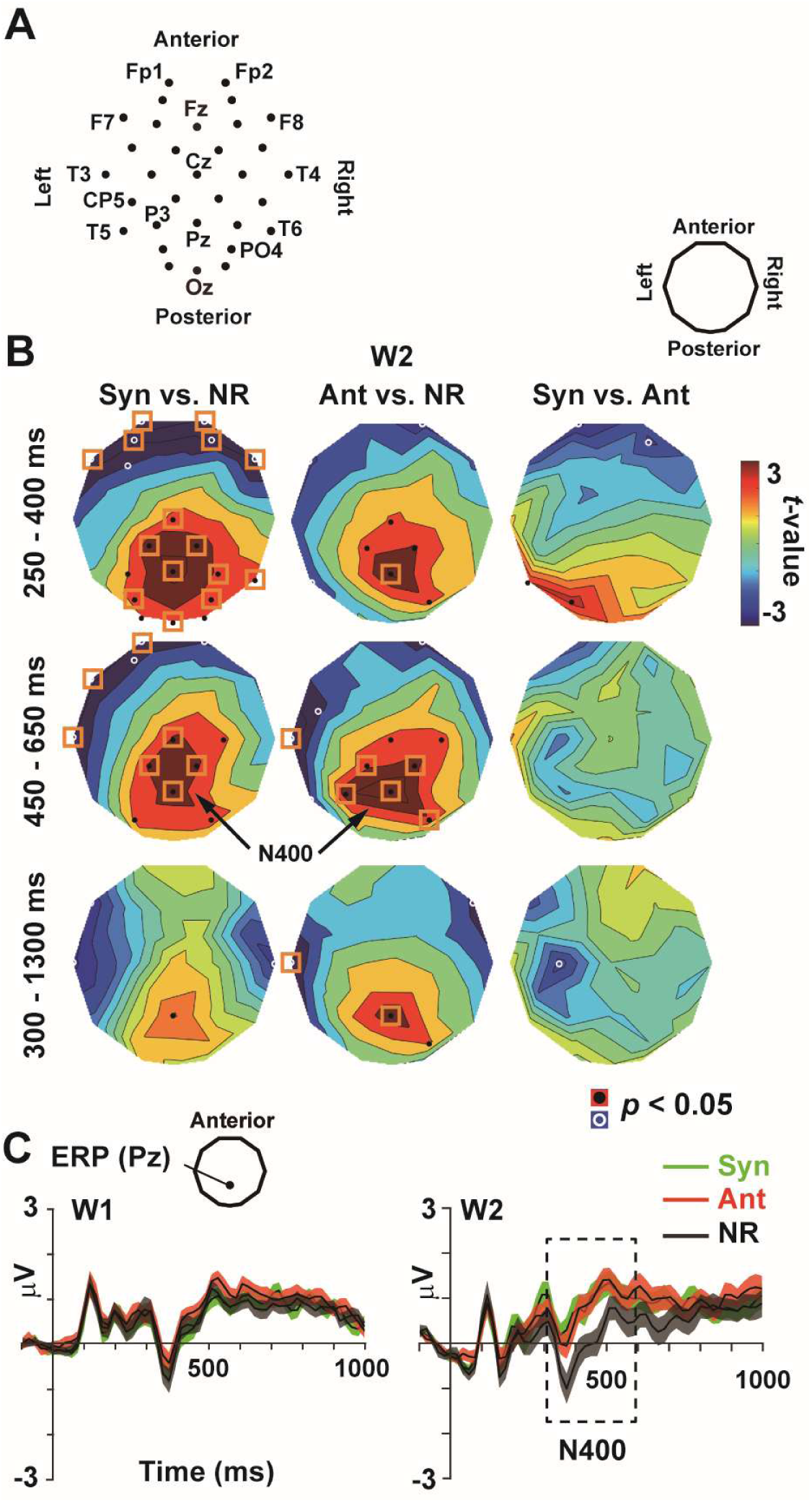
Electroencephalography (EEG). (**A**) Two-dimensional layout of 32 EEG sensors. Sensors over the anterior/posterior brain regions are shown in upward/downward positions. (**B**) *t*-maps of event-related potentials (ERPs) to W2. Mean ERPs at 250 – 400 ms (upper panels), 450 – 650 ms (middle panels), and 300 – 1300 ms (lower panels) were compared across the three conditions. Resultant *t*-values were color-coded over the topographical layout of 32 sensors. Black and white circles denote sensors with positive and negative differences at *p* < 0.05 (uncorrected), while orange rectangles denote significant differences after a correction of multiple comparisons (*p* < 0.05, corrected). No sensor with a significant difference was observed in a direct comparison between Syn vs. Ant (right panel). (**C**) ERP waveforms at Pz in the W1 (left) and W2 (right) periods. Time 0 in a horizontal axis indicates an onset of words. The N400 was observed in the W2 period as a difference between NR (black) and Syn (green)/Ant (red) trials. Background shadings denote standard error (SEs) across participants.

The pre-processing of EEG data were performed by the Brainstorm toolbox (Tadel et al., 2011) implemented in Matlab 2022b. Low-frequency, high-frequency, and fixed-frequency (power-line) noises were suppressed by band-pass (0.5 – 200 Hz) and notch (60, 120, and 180 Hz) filters. All waveforms were then referenced with an average potential over the 32 electrodes. EEG responses to W1 and W2 were segmented (epoch range: −800 to 1900 ms relative to a word onset, **Fig. 1A**) and classified into six conditions (Syn/Ant/NR × W1/W2).

### Analysis of EEG data

I initially computed event-related potentials (ERPs, **Fig. 2B**). EEG waveforms for each of the six conditions were averaged across 84 trials, with a pre-stimulus period (−100 to 0 ms) used as a baseline. Trials were excluded from analyses when a max-min amplitude of EEG waveform was larger than 150 μV (analysis period: −200 to 1300 ms). Numbers of trials that remained after this exclusion (mean ± SD) were 78.38 ± 11.36 (Syn-W1), 72.82 ± 15.34 (Syn-W2), 77.62 ± 12.59 (Ant-W1), 73.44 ± 14.89 (Ant-W2), 77.50 ± 13.37 (NR-W1), and 72.32 ± 16.21 (NR-W2). Based on a previous study (Vaughan et al., 1982), I set two time windows for EPR analysis; 250 – 400 ms and 450 – 650 ms. A mean ERP averaged over 250 – 400 ms was computed for each condition, and statistically compared between conditions in upper panels of **Figure 2B** (see **Statistical procedures** below). The same procedure was performed for a mean ERP at 450 – 650 ms (middle panels).

I then analyzed neural oscillatory signals. To get an overview, an EEG waveform in each trial was first converted into a time × frequency (TF) power spectrum (wavelet transformation). The central frequency of the Morlet wavelet was set at 1 Hz, while its time resolution (FWHM) was 3 s. A baseline correction was also performed using the data in a pre-stimulus period (−200 to 0 ms). The TF spectra averaged across all trials and all participants are shown in **Figure 3**.

**Figure 3.**
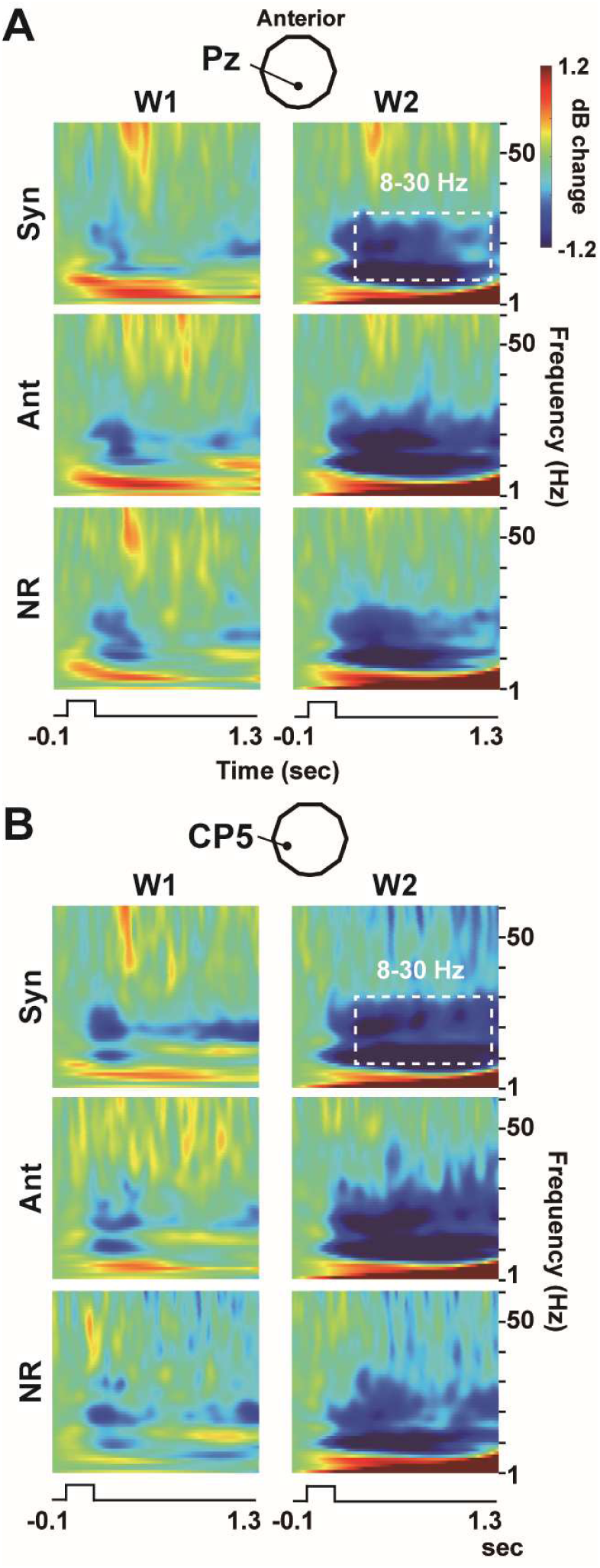
Time-frequency power spectra in six conditions (Syn/Ant/NR × W1/W2). (**A**) Spectra at Pz, an electrode with a distinct N400 response (**Fig. 2**). Decibel power changes from a pre-stimulus period of (−200 to 0 ms) were averaged across all trials and participants. (**B**) Spectra at CP5, an electrode over the left temporo-parietal junction near the Wernicke’s area. At both sensors, sustained and prominent power changes were seen in alpha-to-beta band (8-30 Hz, dotted rectangles), especially in the W2 period (right panels).

Finally, I measured powers and speeds of oscillatory signals in the six conditions (Syn/Ant/NR × W1/W2). An analysis period was set at 300 – 1300 ms after a word onset. Data from 0 to 300 ms were not used, because they might comprise visually-evoked responses to words such as P100 and N170 (Vogel and Machizawa, 2004). An EEG waveform at 300 – 1300 ms was converted into a power spectral density (PSD, frequency resolution: 1 Hz). A power in each frequency band (e.g. delta, theta, … etc.) was obtained by averaging powers of component frequencies belonging to that band. For example, a power in the alpha-to-beta band (see dotted rectangles in **Fig. 3**) was computed by averaging powers of 23 component frequencies from 8 to 30 Hz (mean power). On the other hand, the speed of oscillatory signals was quantified as a central frequency (Klimesch et al., 1993; Samuel et al., 2018) on the same PSD. In case of the alpha-to-beta band, it was obtained by the equation below

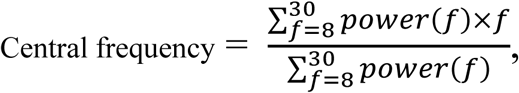

where *f* and *power(f)* indicate the 23 component frequencies (8 – 30 Hz) and their powers, respectively. The mean power and central frequency were computed for each trial and then averaged across all trial in a given condition (e.g. W1 in Syn).

### Statistical procedures

Statistical comparisons of EEG measures were performed by a *t*-test at each of 32 EEG sensors. For example, *t*-maps in left panels in **Figure 2B** show a comparison of ERPs between Syn and NR. In case of upper panel, ERP amplitudes (averaged over 250 – 400 ms) to Syn-W2 of 34 participants were compared with those to NR-W2 through the sensor-wise paired *t*-tests. A problem of multiple comparisons (repetition of *t*-test for 32 times) was resolved by controlling a false discovery rate (FDR). By setting the *q*-value at 0.05 (Li and Noguchi, 2025), a significance threshold was adjusted along with the Benjamini-Hochberg correction (Benjamini and Hochberg, 1995). Sensors with a significant difference after the FDR correction were marked by orange rectangles on each *t*-map.

### Analysis on non-PSD measures

Although I would report the mean power and central frequency as main results (see **Results**), other measures of EEG were used in **Figure S4** (Supplementary Materials), such as Hilbert envelope, instantaneous frequency (IF), and inter-peak interval (IPI). Below are procedures to obtain those alternative (non-PSD) measures.

The Hilbert envelope is a measure for an amplitude of oscillatory signals. A band-pass filter (Butterworth, 8- 30 Hz) was applied to the segmented EEG waveform. An envelope of the filtered waveform (identified with the Hilbert transformation) represented changes in amplitude of the alpha-to-beta rhythm over time. The envelope in each trial was averaged across all trials in a given condition (e.g. Syn-W2). Mean amplitudes at 300 – 1300 ms were compared between conditions through the sensor-wise *t*-tests (**Fig. S4A**, lower panels).

The IF (instantaneous frequency) is a measure for a speed of oscillatory signals (Cohen, 2014). An EEG waveform in alpha-to-beta band was first isolated with a band-pass filter of 8 - 30 Hz. The filtered waveform was then transformed into a phase-angle time series (not the envelope). The IF was obtained as the temporal derivative of this phase-angle timeseries. An oscillation at a higher/lower frequency was indexed by an increased/decreased IF. In each trial, I computed a mean IF at 300 – 1300 ms as a measure for a speed. The IF averaged over trials were administered to the paired *t*-tests between conditions (**Fig. S4B**, upper panels).

The IPI (inter-peak interval) is also a measure for a speed (Noguchi and Kakigi, 2020). As performed in the IF analysis, a band-pass filter of 8 – 30 Hz was applied to the segmented EEG waveform. Each peak of the filtered waveform was identified, and a time interval between contiguous peaks was defined as an IPI. A mean length of all IPIs identified at 300 – 1300 ms provided a measure for a speed, because a slow/fast oscillation resulted in a longer/shorter mean IPI. The mean IPIs pooled across all trials were compared between conditions on *t*-maps in lower panels of **Fig. S4B**.

### Experiment 2

In Experiment 2, I investigated EEG signals for the processing of thematic and taxonomic relationships. A new set of 336 words was made to compare three conditions: Them, Tax, and NR (**Fig. 6A**). The W1 and W2 in Them trials had a relation based on co-occurrence in events or situations (e.g. “desert” and “camel”). Those in Tax trials had common features and thus belonged to the same semantic category (e.g. “rabbit” and “camel”, sharing the features as animals). The shared categorical information was variable across 84 trials, such as foods, beverages, tools, furniture, clothes, places, vehicles, animals, occupations, and body parts, etc. Word pairs in NR trials had neither relationship (e.g. “chain” and “camel”). As Experiment 1, the three types of trials shared the same W2 (“camel”) to control across-condition variabilities in visual inputs. Participants answered a relation between W1 and W2 by choosing one of three options (thematic, taxonomic, or non-related).

All 336 words in Experiment 2 were written in 1-6 kana and kanji letters. Mean numbers of letters and SDs across 84 words were 2.98 ± 1.14 (W1 in Them), 2.86 ± 1.02 (W1 in Tax), 2.80 ± 1.04 (W1 in NR), and 2.80 ± 1.04 (W2). No main effect was observed (*F*(3,332) = 0.53, *p* = 0.66, *η*^2^ = 0.005). Mean (± SD) numbers of moras were 3.39 ± 0.82 (W1 in Them), 3.31 ± 0.76 (W1 in Tax), 3.20 ± 0.79 (W1 in NR), and 3.20 ± 0.79 (W2). No main effect was observed (*F*(3,332) = 1.15, *p* = 0.33, *η*^2^ = 0.010). Finally, cosine similarities between W1 and W2 (mean ± SD) were 0.408 ± 0.102 (Them), 0.404 ± 0.094 (Tax), and 0.198 ± 0.067 (NR). A one-way ANOVA indicated a significant main effect (*F*(2,249) = 153.77, *p* < 0.001, *η*^2^ = 0.553). Post-hoc comparisons showed significant differences in Them vs. NR (corrected *p* < 0.001) and Tax vs. NR (corrected *p* < 0.001), but not in Them vs. Tax (uncorrected *p* = 0.77). Other details were identical to Experiment 1.

## Results

### Behavioral data in Experiment 1 (synonyms vs. antonyms)

Mean (± SE) accuracy of 34 participants in the Sem task was 99.14 ± 0.14%, indicating that they correctly understood the meaning of 336 words. In the Rel task, hit rates in three types of trials were 91.90 ± 0.68 % (Syn), 93.88 ± 0.78 % (Ant), and 98.44 ± 0.21 % (NR), while false-alarm (FA) rates were 1.24 ± 0.18 % (Syn), 1.41 ± 0.13 % (Ant), and 5.29 ± 0.54 % (NR). The d-primes (*d*’s) computed from the hit and FA rates were 3.82 ± 0.09 (Syn), 3.93 ± 0.10 (Ant), and 3.95 ± 0.08 (NR). A one-way ANOVA with the Greenhouse-Geisser correction yielded no main effect (*F*(2,66) = 2.27, *p* = 0.11, *η*^2^ = 0.064). These results suggested that task difficulties were balanced across the three types of trials.

### ERP data

**Figure 2B** shows results of ERP analysis. Based on a previous study (Vaughan et al., 1982), I set two time periods for ERP analysis; 250 – 400 ms and 450 – 650 ms. In upper panels, mean ERPs at 250 – 400 ms were compared between Syn-W2 vs. NR-W2 (left), Ant-W2 vs. NR-W2 (middle), and Syn-W2 vs. Ant-W2 (right). The *t*-values of the sensor-wise *t*-tests are color-coded over a two-dimensional layout of 32 sensors. The same analysis was performed with the time window set at 450 – 650 ms (middle panels).

Consistent with previous studies (Roehm et al., 2007; Lau et al., 2008), clear N400 responses to unrelated (NR) compared to related (Syn and Ant) pairs of words were observed over the parietal sensors. No significant difference, however, was observed in a direct comparison of Syn-W2 vs. Ant-W2 (right panels). Results were unchanged when an analysis period was extended to 300 – 1300 ms (lower panels in **Fig. 2B**). These data indicated that ERPs did *not* differentiate synonyms from antonyms, although they discriminated related (Syn and Ant) from unrelated (NR) pairs of words.

### Power in oscillatory signals

**Figure 3A** shows TF (time-frequency) spectra at Pz where clear N400 response was observed. In all conditions (Syn, Ant, and NR), prominent and sustained power changes (decreases) were seen in alpha-to-beta band (8 – 30 Hz, marked by a dotted rectangle). Those changes in alpha-to-beta power were distinct in the W2 period (right panels), suggesting that the changes reflected neural activity related to a judgment of semantic relationships. Similar results were observed at CP5 (**Fig. 3B**), an electrode over the left temporo-parietal junction (Sowden et al., 2015) near the Wernicke’s area. These results were consistent with previous studies reporting an involvement of alpha and beta rhythms in semantic processing (Hall et al., 2025; Rempe et al., 2025) and verbal working memory (Meltzer et al., 2017; Proskovec et al., 2019; Noguchi, 2024; Diedrich et al., 2025). Therefore, I would report oscillatory signals of this band in main texts. Data in other frequency bands (delta, theta gamma, and high-gamma) are shown in **Figure S1 and S2** of **Supplementary Materials**.

Results of statistical analyses on oscillatory powers are displayed in **Figure 4A**. An EEG waveform at 300 – 1300 ms was converted into a power-spectral density (PSD), and a mean power on the PSD (8- 30 Hz) was compared between conditions through the sensor-wise *t*-tests. A *t*-map in the left panel showed a reduction of the alpha-to-beta power in Syn-W2 compared to NR-W2. A similar reduction of power was also seen in Ant-W2 vs. NR-W2 (middle panel). A direct comparison between Syn-W2 vs. Ant-W2, however, showed no significant difference at any sensors (right panel).

**Figure 4.**
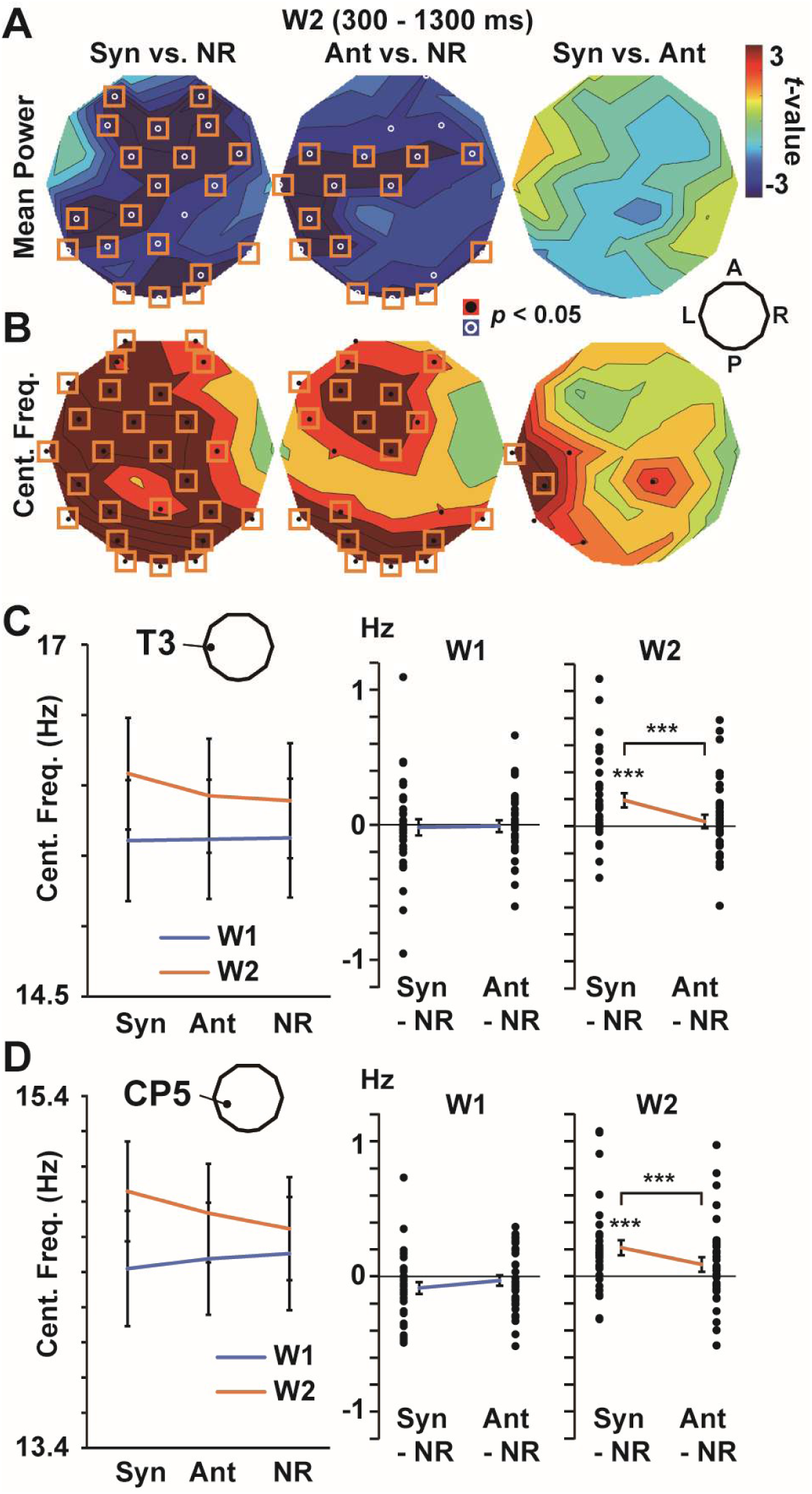
Statistical comparisons of neural oscillatory signals (8 – 30 Hz). (**A**) *t*-maps of mean power. (**B**) *t*-maps of central frequency. Both measures were obtained from a power-spectral density (PSD) of the EEG waveform at 300 – 1300 ms relative to W2 onset. A direct comparison of Syn vs. Ant (right panels) showed significant differences at left temporal sensors (T3 and CP5) in central frequency, but not in mean power. (**C**) Changes in central frequency in the six conditions at T3, a sensor over the anterior temporal cortex. Differences from a control condition (Syn – NR and Ant - NR) are shown in middle (W1) and right (W2) panels. Individual data of 34 participants are denoted by black dots, while error bar indicate SEs. (**D**) Change in central frequency at CP5. \*\*\**p* < 0.001.

This lack of difference in Syn-W2 vs. Ant-W2 was not caused by the present approach in which powers in alpha and beta bands were jointly analyzed. I showed in **Figure S3** the *t*-maps where mean powers in alpha (8 – 13 Hz) and beta (13- 30 Hz) bands were separately analyzed. No significant difference between Syn and Ant was observed in alpha (upper right panel) or beta (upper left panel) power.

### Speeds in oscillatory signals

I then analyzed the central frequency, a measure for a speed of neural oscillations. A comparison of Syn-W2 vs. NR-W2 (**Fig. 4B**, left panel) showed a significant increase (Syn > NR) in central frequency over widespread regions. An increase was also seen in Ant-W2 vs. NR-W2 (middle), although the sensors over the temporal cortex showed no difference. Consequently, a comparison of Syn-W2 vs. Ant-W2 (right) revealed significant increases (Syn > Ant, corrected *p* < 0.05) at left temporal sensors, such as T3 and CP5. I displayed in **Figure 4C** the central frequencies at T3 in three conditions. Differential values from a control condition, such as Syn – NR and Ant – NR, were provided in right panels. In the W2 period, central frequencies in Syn were significantly higher than that of NR (*t*(33) = 3.63, *p* = 0.00095, Cohen’s *d* = 0.08). The difference between Syn vs. Ant was also significant (*t*(33) = 3.95, *p* = 0.0004, *d* = 0.07). Similar results were seen at CP5 (**Fig. 4D**, Syn vs. NR: *t*(33) = 3.84, *p* = 0.0005, *d* = 0.13, Syn vs. Ant: *t*(33) = 3.67, *p* = 0.0009, *d* = 0.08).

### Results of non-PSD measures for oscillation amplitude and speed

PSD measures (mean power and central frequency) indicated that neural signals differentiating Syn from Ant were embedded in a temporal domain of EEG data. I tested a robustness of these results using non-PSD measures. In **Figure S4A**, amplitudes of oscillatory signals (8 – 30 Hz) were quantified through the wavelet analysis (upper panels) and Hilbert transformation (lower panels). No sensor showed a difference between Syn-W2 vs. Ant-W2 even at an uncorrected level of *p* < 0.05. **Figure S4B** shows *t*-maps for measures of speed, such as instantaneous frequency (IF, upper panels) and inter-peak interval (IPI, lower panels). Note that a faster neural oscillation is indexed by higher IF and shorter IPI. Both measures indicated an acceleration of brain rhythm in Syn-W2 compared to Ant-W2 over the left temporal cortex (right panels). Those results provided converging evidence that information about conceptual similarity was mainly encoded in temporal measures of neural activity.

### Experiment 2 (thematic vs. taxonomic relationships)

Data in Experiment 1 showed faster neural responses induced by synonyms than antonyms. These results are consistent with a classic model of neural network in which related semantic concepts are connected with each other (Quillian, 1967; Collins and Loftus, 1975). A spread of neural activation from W1 temporally facilitated the processing of W2 in Syn trial, because W1 (e.g. “progress”) and W2 (“advance”) were located nearby in the network (**Fig. 5A**). No facilitation was induced by an antonym pair of words, since their opposite meanings were characterized by a longer distance.

**Figure 5.**
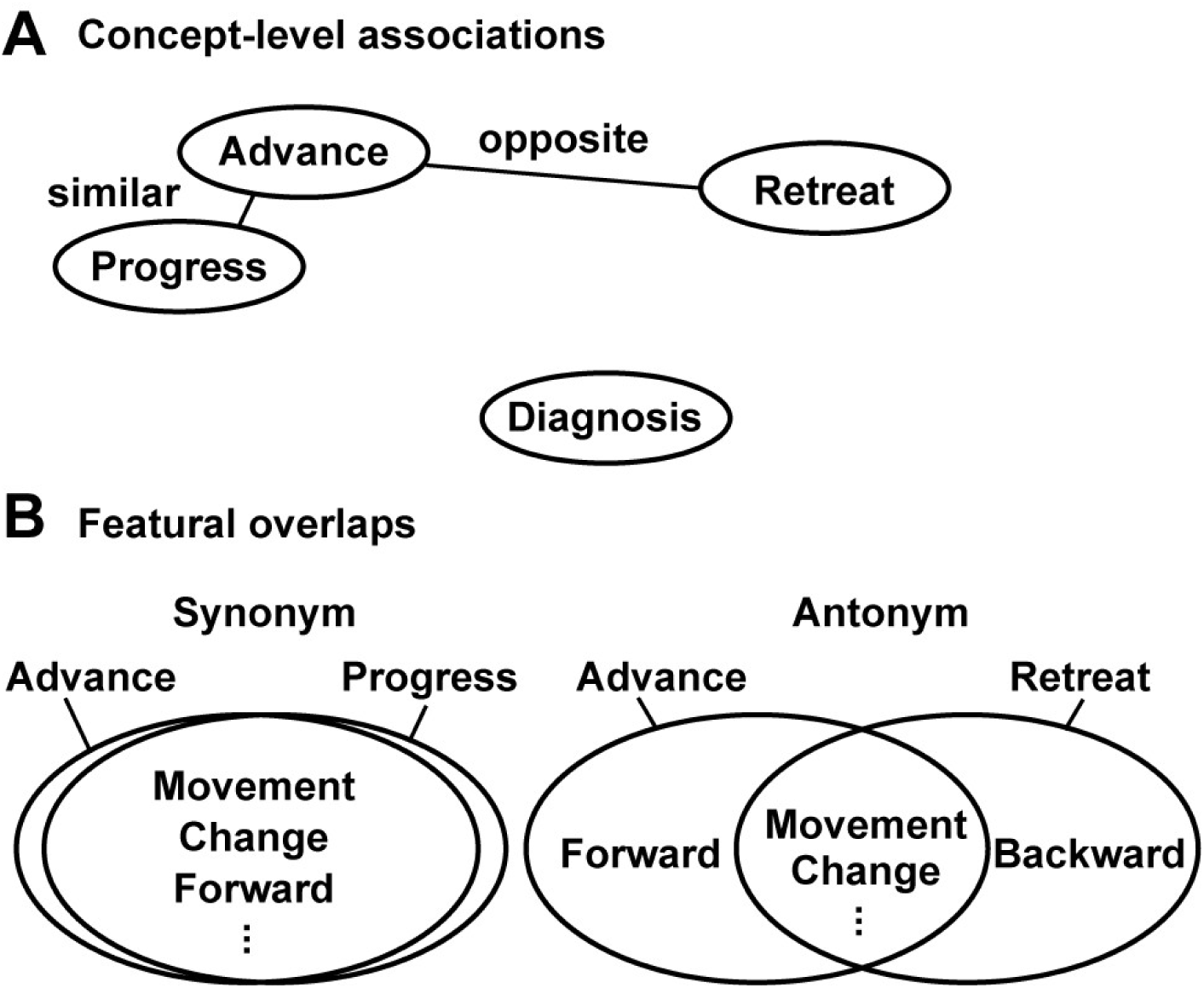
Two possibilities accounting for the results of Experiment 1. (**A**) An account based on semantic (conceptual) network in the brain. A synonym pair of words are connected and located nearby because of their conceptual similarity, while words in antonymy are characterized by a longer distance because of their opposite meanings. This difference in inter-conceptual distances induced a spread of neural activity from W1 to W2 selective to Syn trials, resulting in an increase of central frequency (Syn-W2 > Ant-W2, **Fig. 4C**). (**B**) An account based on featural overlaps. While a synonym pair shared a nearly-identical set of semantic features, an antonym pair differed in a critical feature (e.g. forward vs. backward in case of “advance” and “retreat”). A total degree of featural overlaps was larger in Syn than Ant trials, which produced an acceleration (temporal facilitation) of the semantic processing in Syn-W2.

Another possibility that can explain the results, however, is a difference in overlaps of semantic features (**Fig. 5B**). While a synonym pair of words have a virtually-identical set of features, an antonym is two words that are similar in all but one aspect (Willners, 2001; Scheible et al., 2013; Kliegr and Zamazal, 2018). A total degree of featural overlaps is thus larger in synonym than antonym pairs, which might produce the higher central frequency in Syn than Ant trials (**Fig. 4B**).

To discern those two possibilities (conceptual closeness vs. featural overlap), I conducted Experiment 2 in which EEG signals were compared between thematically- and taxonomically-related pairs of words (**Fig. 6A**). The W1 and W2 in Them trials had a relation based on co-occurrence in scenes or events (e.g. “desert” and “camel”), while those in Tax trials had common categorical features (e.g. “rabbit” and “camel”). Participants judged a relation between W1 and W2 by selecting one of three options (thematic, taxonomic, or non-related). Note that words in Tax trials were characterized by shared features between W1 and W2, while those in Them trials shared no features but were located close at a concept level. If the higher central frequency in Syn than Ant trials (**Fig. 4B**) reflected a degree of featural (elemental) overlaps (**Fig. 5B**), the same region would show a higher central frequency in Tax than Them trials. In contrast, the increased frequency in Syn trials reflected conceptual (holistic) closeness between W1 and W2, the higher frequency might be expected in Them trials.

**Figure 6.**
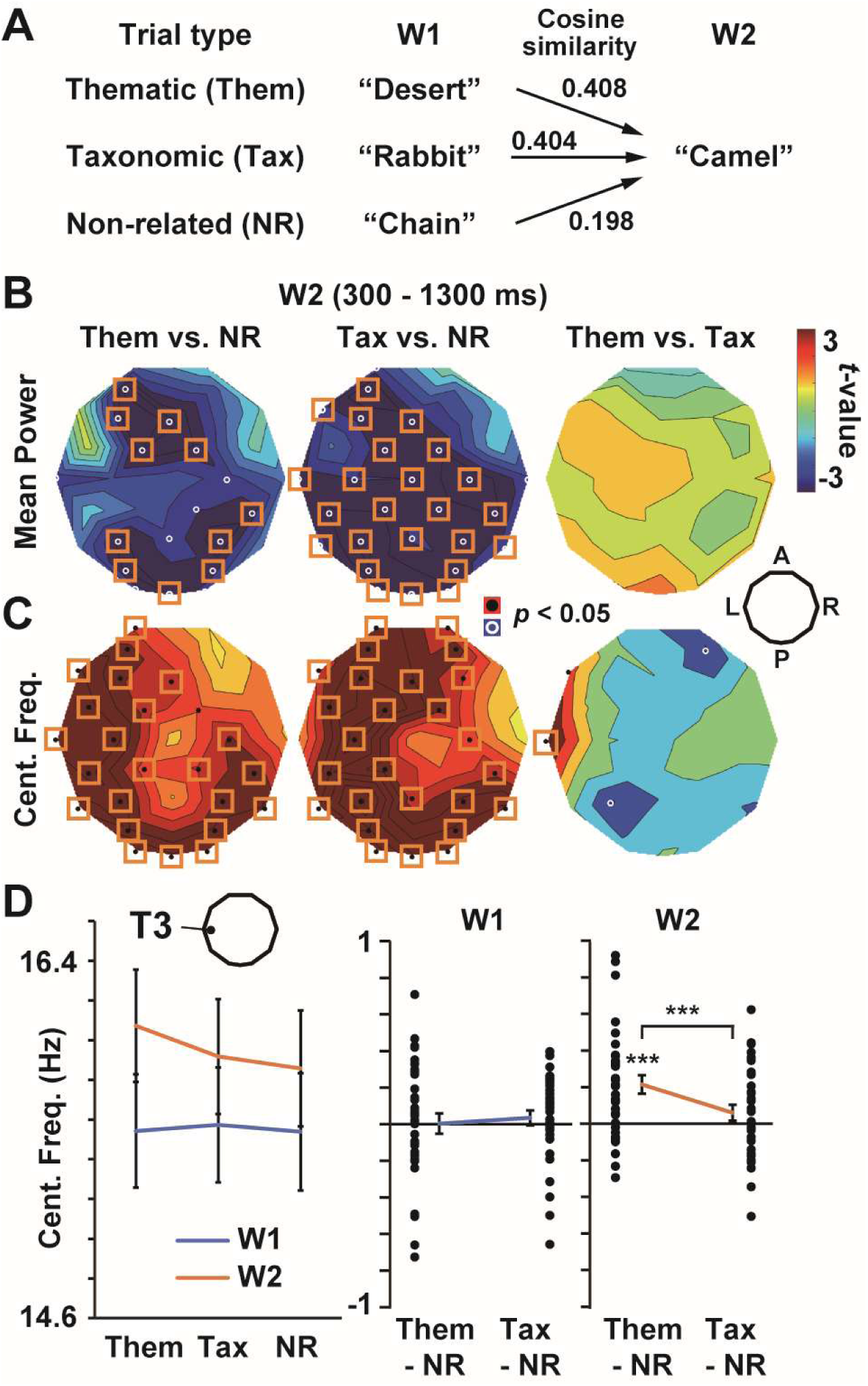
Experiment 2. (**A**) Structures of the Rel task. W1 and W2 were thematically related, taxonomically related, or unrelated in Them, Tax, and NR trials, respectively. The taxonomic relations were based on shared categorical features (e.g. “rabbit” and “camel”, sharing the features as animals), while thematic relation were not. If the changes of central frequency in Experiment 1 reflected a degree of featural overlaps (**Fig. 5B**), oscillatory signals over the left temporal regions would be accelerated in Tax rather than Them trials. (**B**) *t*-maps of mean power. (**C**) *t*-maps of central frequency. An increase in central frequency was seen in Them than Tax trials at T3 (right panel), indicating that changes in brain rhythm reflected the inter-conceptual closeness (memory-based association) between W1 and W2. (**D**) Individual data of central frequency at T3. \*\*\**p* < 0.001.

Behavioral data showed that 34 participants correctly understood all (336) words, and that task difficulties were balanced across the three types of trials. Mean ± SE accuracy of 34 participants in the Sem task was 98. 98 ± 0.18 %. The d-primes (*d*’s) in the Rel task were 3.87 ± 0.09 (Them), 4.01 ± 0.11 (Tax), and 3.88 ± 0.07 (NR), with a one-way ANOVA indicating no main effect (*F*(1.71, 56.26) = 2.32, *p* = 0.12, *η*^2^ = 0.066).

Results in EEG data (alpha-to-beta band) are shown in **Figure 6B-D** (data in other frequency bands are shown in **Fig. S5** and **Fig. S6**). Mean powers on PSDs (**Fig. 6B**) to W2 showed significant differences between Them vs. NR (left panel) and Tax vs. NR (middle), but not in Them vs. Tax (right). On the other hand, central frequencies (**Fig. 6C**) were significantly higher in Them than Tax trials at an anterior temporal electrode (T3, right panel). Individual data of Them – NR and Tax – NR (**Fig. 6D**) indicated a higher central frequency in Them than NR in the W2 period (*t*(33) = 4.30, *p* = 0.0001, *d* = 0.13). The difference between Them vs. Tax was also significant (*t*(33) = 4.32, *p* = 0.0001, *d* = 0.09).

## Discussion

I presently measured EEG signals in response to various semantic relations of words. As the first analysis, I compared ERPs across three conditions in Experiment 1; Syn, Ant, and NR (**Fig. 2**). It has been controversial whether ERPs can differentiate synonyms from antonyms; Vaughan et al. (1982) reported different patterns of ERPs between synonyms and antonyms (Vaughan et al., 1982), although no such differences were observed in Thatcher (1977) and Goto et al. (1979) (Thatcher, 1977; Goto et al., 1979). Vaughan et al. (1982) measured EEG at CPz from five subjects and reported a “biphasic” ERP response; evoked potentials to synonyms were more positive than those to antonyms at 250 – 400 ms (Syn > Ant), while this relationship was reversed at 450 – 650 ms (Syn < Ant). I thus compared mean ERPs at those periods in upper and middle panels in **Figure 2B**. No significant difference, however, was seen in Syn vs. Ant (right panels) at either period. It is thus hard to discern synonyms from antonyms with ERPs, especially when relatedness of W1 and W2 (cosine similarity) was strictly balanced between Syn and Ant conditions as in the present study (**Fig. 1B**).

I then analyzed neural oscillatory signals, focusing on a temporal measure. In Experiment 1, I observed an increase of central frequency in Syn vs. NR and Ant vs. NR (**Fig. 4B**. left and middle panels) in widespread regions over the occipital, parietal, and medial frontal cortices. A direct comparison between synonym and anonym, on the other hand, showed an increase localized to the left temporal areas (right panel). These results suggest an integrated model of the “core-network” view (Binder et al., 2009; Ralph et al., 2017; Fedorenko et al., 2024) and the “distributed” view (Huth et al., 2016; Anderson et al., 2017) in previous studies. The widespread changes common to synonyms and antonyms (occipito-parietal and medial-frontal) indicate that those regions reacted to relatedness of W1 and W2 based on an overlap of their semantic features. This is consistent with a previous view that semantic information is distributed over broad regions of the human brain (Huth et al., 2016; Anderson et al., 2017). In contrast, the selective increase in central frequency to Syn (right panel) indicates that the left temporal region encoded a W1-W2 similarity at a concept level. This highlights a special role of this area, consistent with the “core-network” theory (Binder et al., 2009; Ralph et al., 2017; Fedorenko et al., 2024). In short, data in Experiment 1 suggest a gradual change of semantic processing from feature-based to concept-based mechanisms represented in posterior-anterior axis in the left hemisphere (**Fig. 7**). This separation between the two mechanisms might enable us to understand a “related but dissimilar” relationship of words typically seen in antonym pairs (Hill et al., 2015).

**Figure 7.**
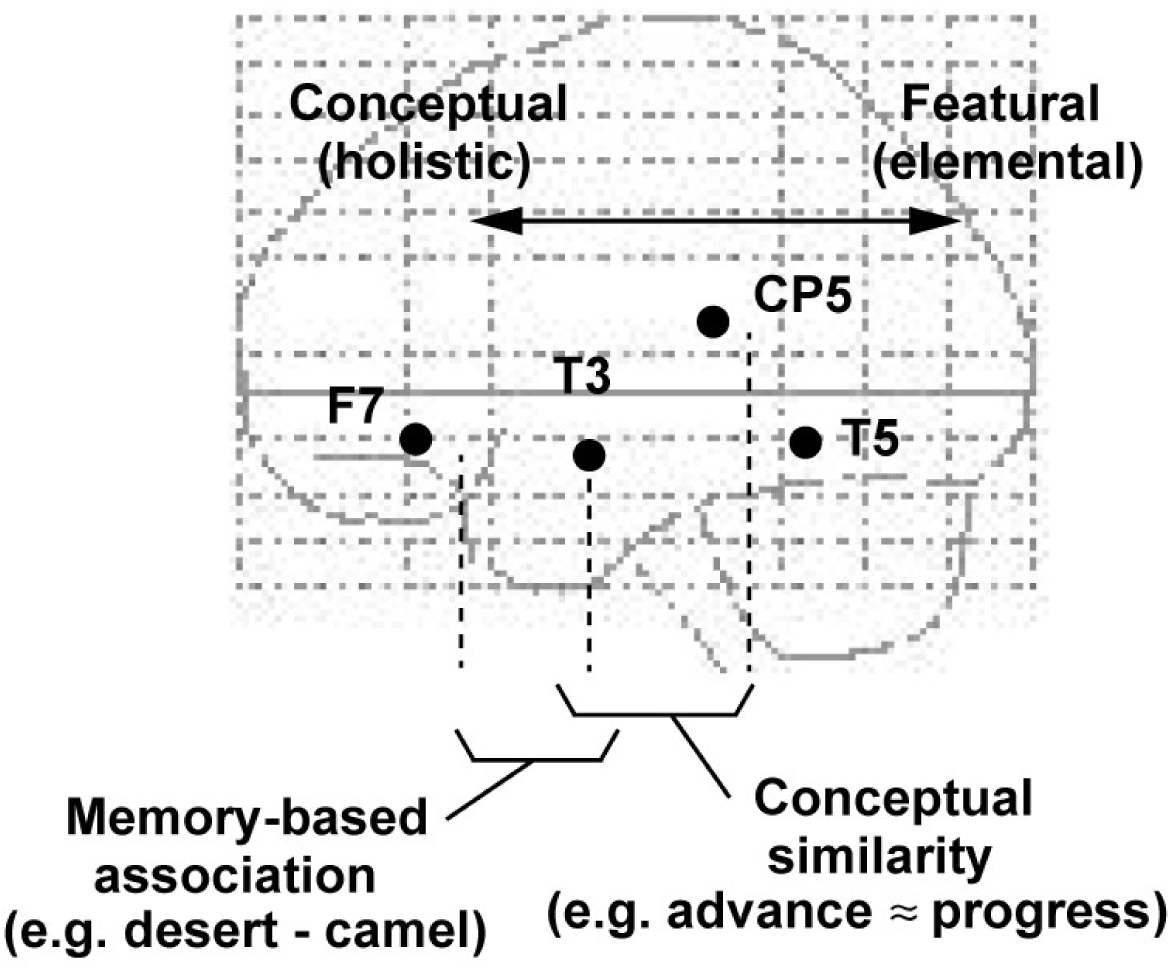
A model suggested by the present data. Experiment 1 showed a temporal facilitation of EEG signals in Syn than Ant trials at T3 and CP5, indicating the neural processing in this region based on conceptual similarity or identity. Experiment 2 further showed that an anterior part of this region (T3) also encoded inter-conceptual closeness based on declarative memories (thematic relation). In contrast, a posterior part of the brain (occipital and parietal areas) reacted to both Syn and Ant in Experiment 1 (**Fig. 4B**, left and middle panels). The central frequency over the left parietal region tended to be higher in Tax than Them trials, although this difference was not significant after the correction (**Fig. 6C**, right). These results suggest a dominance of feature-based processing in the posterior regions.

In Experiment 2, I further investigated neural activity for the processing of thematic vs. taxonomic relationships, a matter of intense debate in a previous literature (Kuchinke et al., 2009; Sachs et al., 2011; de Zubicaray et al., 2013; Jackson et al., 2015; Lewis et al., 2015; Kumar, 2018; Xu et al., 2018; Chou et al., 2019; Carota et al., 2021; Thye et al., 2021; Blackett et al., 2022; Fernandino et al., 2022; Fu et al., 2023; Zhang et al., 2023; Adezati et al., 2024; Liu et al., 2024; Riccardi et al., 2024; Stochel and Sandberg, 2025). Results revealed an increase in central frequency specific to the Them trials at T3, an electrode over the anterior temporal cortex (ATL). A comparison between Tax-W2 and NR-W2 (**Fig. 6C**, middle panel) showed no significant change at this sensor (*t*(33) = 1.37, *p* = 0.18, *d* = 0.04). These results reinforce the view in Experiment 1 that the ATL reacted to conceptual (holistic) closeness between words, not their featural (elemental) overlaps. We know through our experiences that W1 and W2 in Them trials (e.g. “camel” and “desert”) co-occur in the same scene or event, although they belong to different semantic categories. An increased central frequency in the ATL would reflect this memory-based association (closeness) in Them pairs that cannot be detected by featural (categorical) overlaps.

It is surprising that the ATL showed a selective response to Them trials, because the dual-hub theory (Schwartz et al., 2011) assumes a role of this region in processing taxonomic relations. Several recent studies, however, reported an involvement of ATL in thematic processing (Blackett et al., 2022; Zhang et al., 2023; Riccardi et al., 2024). Furthermore, the present results are consistent with a classic view of neuropsychology that knowledge in declarative (episodic and semantic) memories is stored in the ATL (Kapur et al., 1992; Mummery et al., 2000; Nestor et al., 2006; Herlin et al., 2021; Setton et al., 2022). Given that many thematic relations are grounded in declarative memories (Li et al., 2011; Zhang et al., 2023), the present results provided converging evidence for the ATL as a center of conceptual (verbal) knowledge, filling a gap between memory and linguistic research.

Importantly, all results described above were obtained through an analysis of central frequency. No change in oscillation amplitude/power was observed in critical comparisons (Syn vs. Ant and Them vs. Tax). These results indicate an importance of temporal measures in analyzing neural activity. Given a tight coupling of oscillation amplitude with hemodynamic signals (Murta et al., 2015; Bondi et al., 2025; Xavier et al., 2025), the present results would explain why previous studies using hemodynamic measures have provided mixed results on brain regions for thematic and taxonomic processing (Kuchinke et al., 2009; Sachs et al., 2011; de Zubicaray et al., 2013; Jackson et al., 2015; Kumar, 2018; Xu et al., 2018; Chou et al., 2019; Carota et al., 2021; Fernandino et al., 2022; Fu et al., 2023; Liu et al., 2024; Stochel and Sandberg, 2025).

What type of neural processing underlies the increase in central frequency observed in the present study? One possibility is a priming effect in neural signals. Mounting evidence from behavioral studies (Lucas, 2000; Hutchison, 2003) supported a network model of semantic memory in which related concepts are connected with each other (Quillian, 1967). It is highly possible that W1 and W2 in synonym trials (e.g. “progress” and “advance”) were directly connected in this network (**Fig. 5A**), because of their high conceptual similarity. A serial presentation of W1 and W2 thus temporally facilitated the semantic processing of W2 through the spreading of neural activation from W1(Collins and Loftus, 1975), resulting in an acceleration of EEG signals (Noguchi et al., 2004). A major limitation of the present study is a poor spatial resolution of scalp EEG. An approach with a higher spatial resolution, such as electrocorticography, is helpful to further reveal mechanisms of the present effect.

## Acknowledgments

This work was supported by grants from the Japan Society for the Promotion of Science (KAKENHI: 25K06896), from The Fukuhara Fund for Applied Psychoeducation Research (Tokyo, Japan), and from The Mitsubishi Foundation (ID: 202520025) to Y.N. I thank Keigo Furusho, Yu Fujimoto, and Taeko Kaneda for their support. All data supporting the findings of this study are available from Y.N. upon reasonable request.

## Author contributions

Y.N. developed the study concept, prepared stimuli, collected data, performed data analysis, and wrote the paper.

## Supplementary Materials

**Figure S1.**
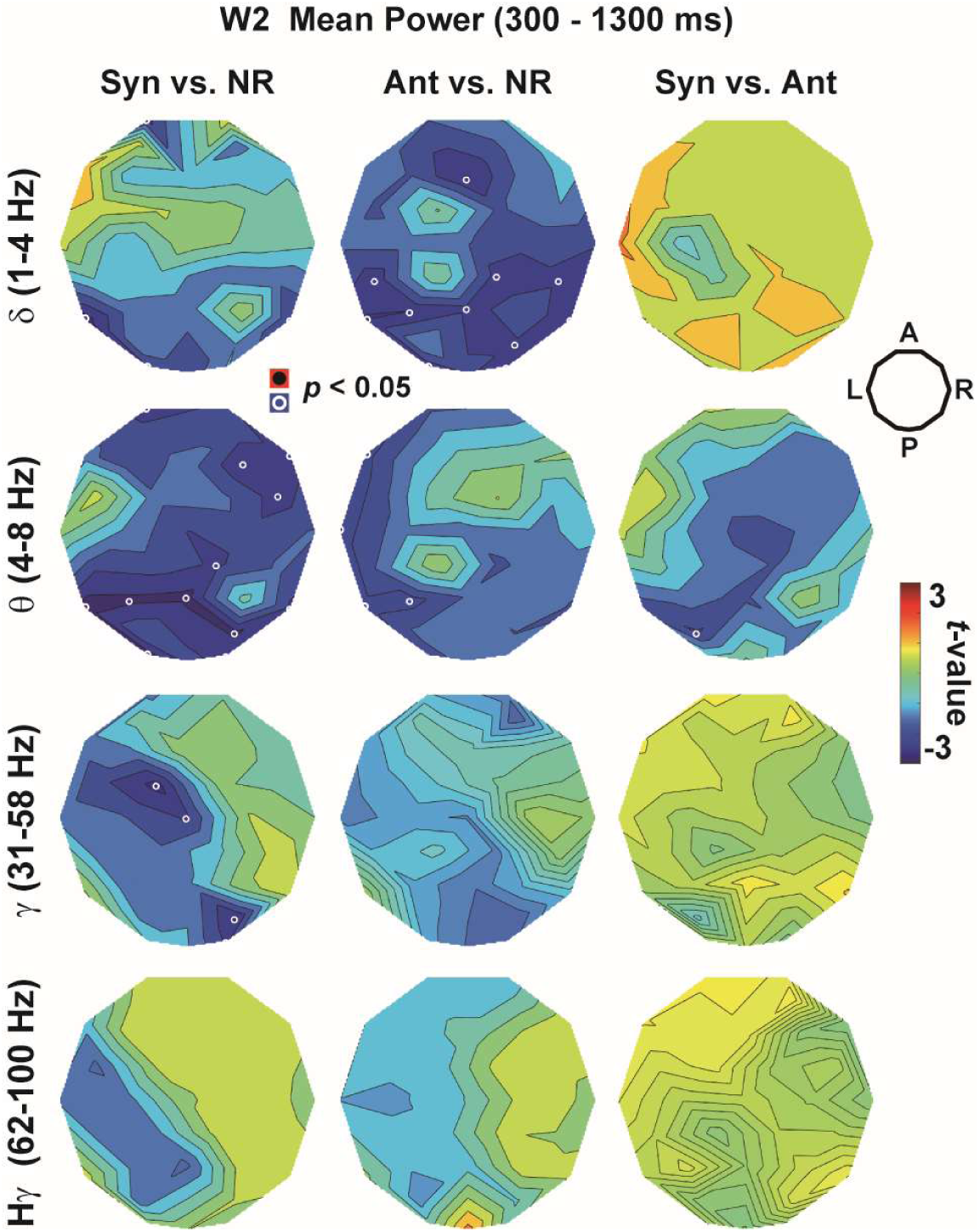
The *t*-maps of mean power in Experiment 1. Data in delta (1 – 4 Hz), theta (4 – 8 Hz), gamma (31 – 58 Hz), and high-gamma (62 – 100 Hz) bands are shown. No sensor showed significant difference after the correction of multiple comparisons.

**Figure S2.**
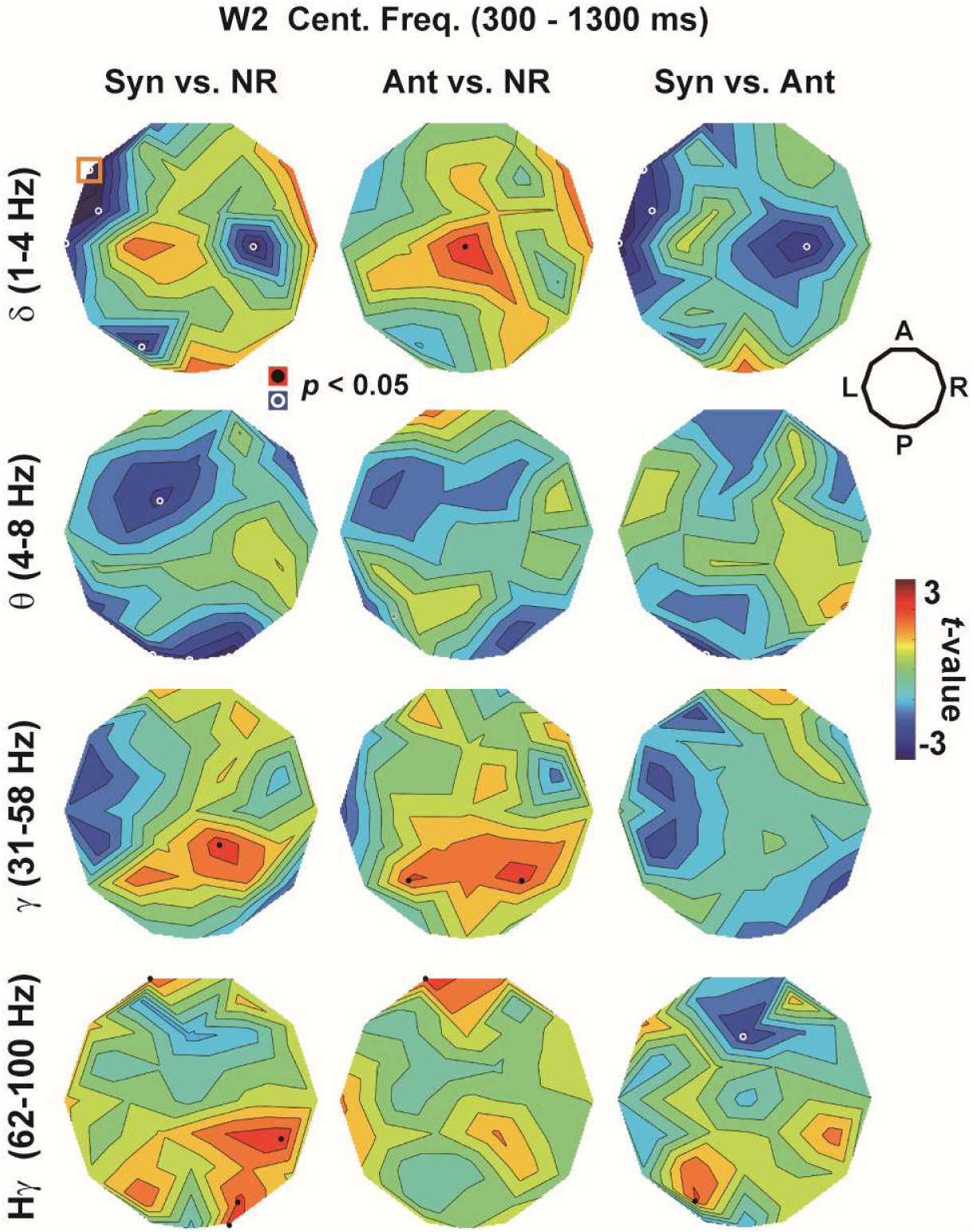
The *t*-maps of central frequency in Experiment 1 in delta, theta, gamma, and high-gamma bands. No sensor showed a significant difference, except for F7 (Syn vs. NR) in delta band (denoted by an orange rectangle, *p* < 0.05, corrected).

**Figure S3.**
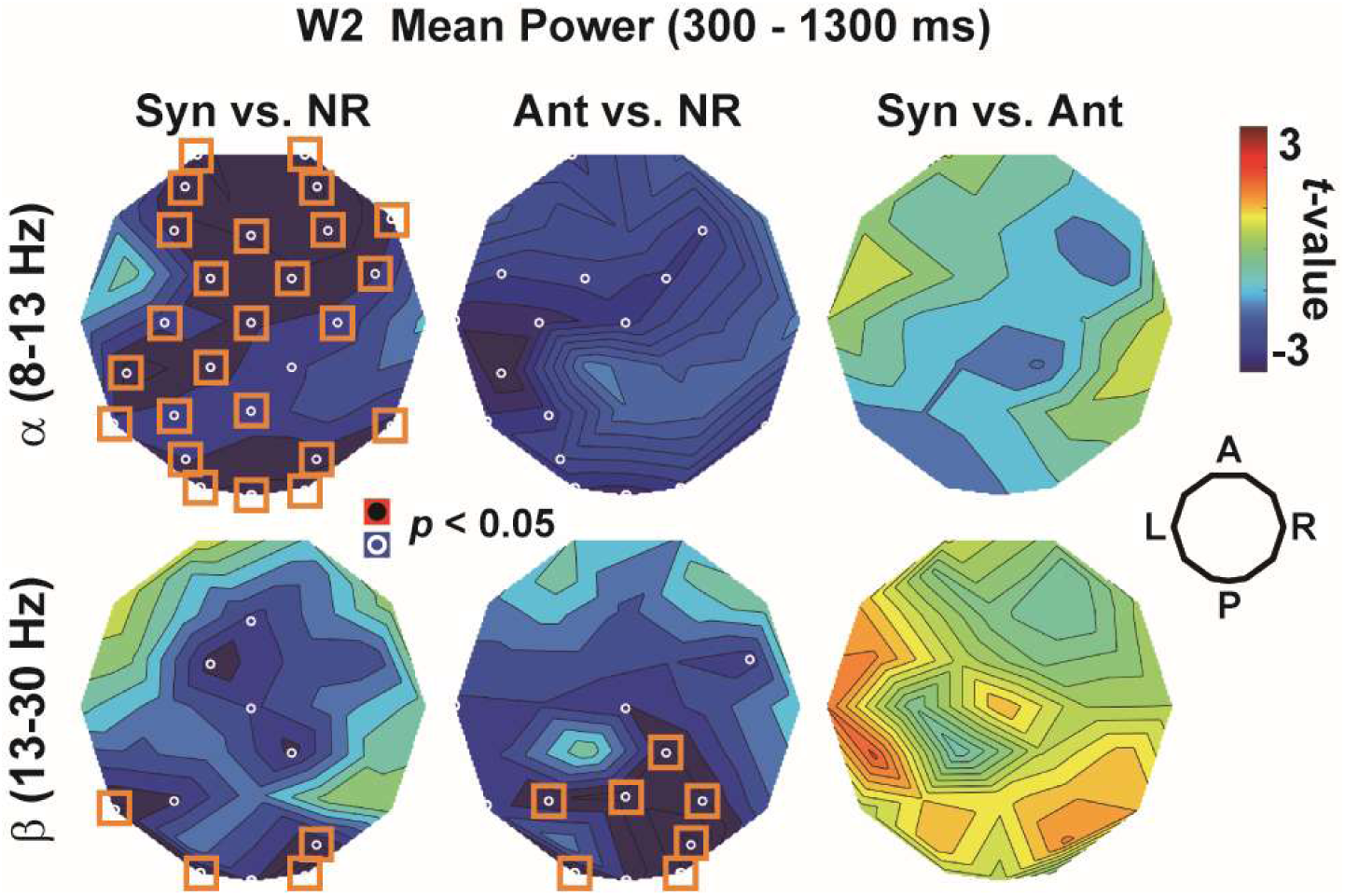
The *t*-maps of mean power in Experiment 1. Data of alpha (8 – 13 Hz) and beta (13 – 30 Hz) bands are separately shown. Although significant decreases in alpha and beta powers were seen in Syn vs. NR (left panels) and Ant vs. NR (middle panels), no sensor showed a significant difference in Syn vs. Ant (right panels).

**Figure S4.**
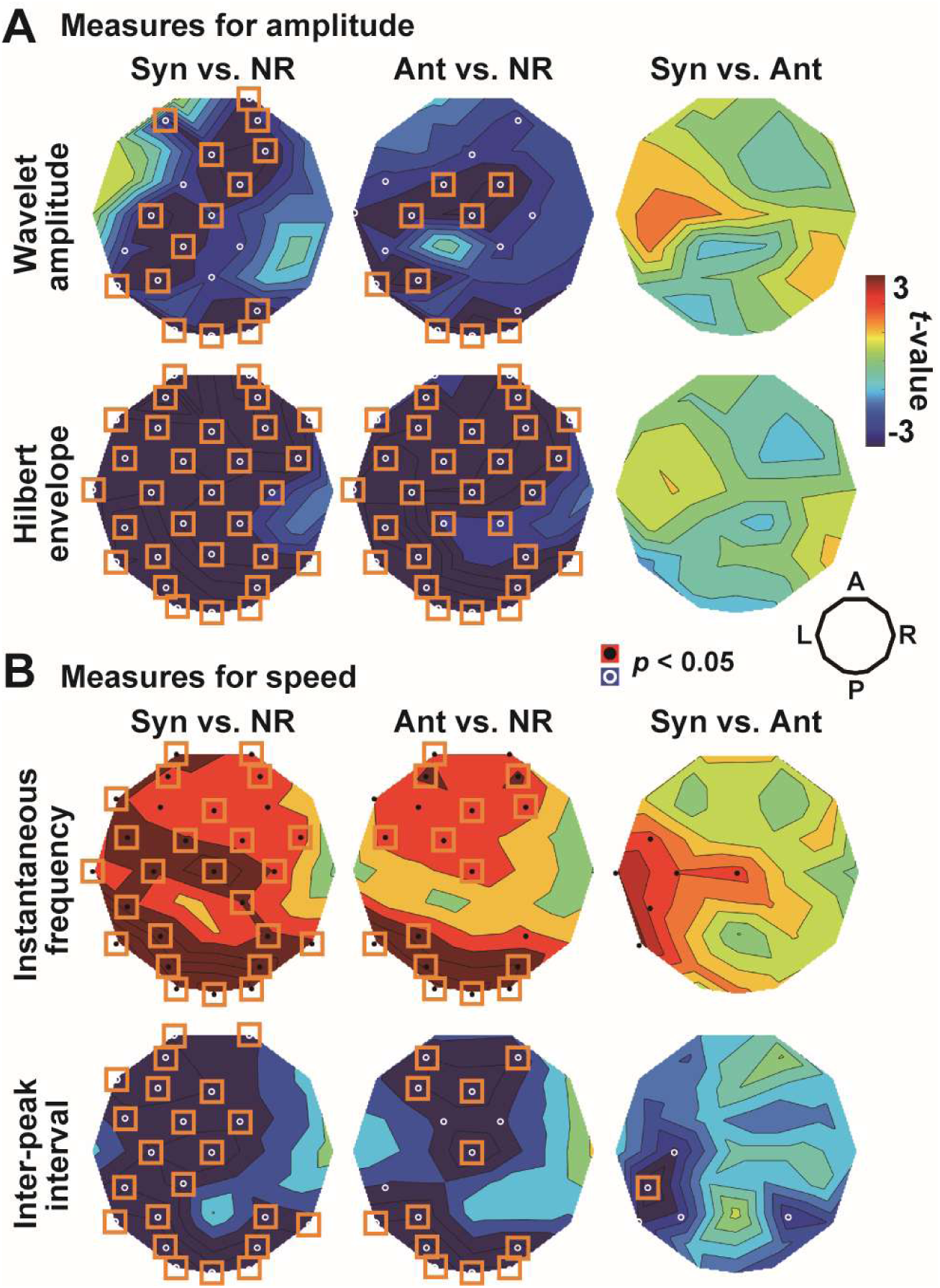
Results of non-PSD measures in Experiment 1. (**A**) *t*-maps of the wavelet amplitudes (upper panels) and Hilbert envelopes (lower panels), both of which are measures for amplitude of neural oscillatory signals. (**B**) *t*-maps of measures for speed; instantaneous frequency (IF, upper panels) and inter-peak interval (IPI, lower panels). Note that a higher speed of oscillation is indexed by a higher IF and shorter IPI. Differences between Syn and Ant (right panels) at left temporal sensors were more clearly seen in the speed than amplitude measures, consistent with results in main texts.

**Figure S5.**
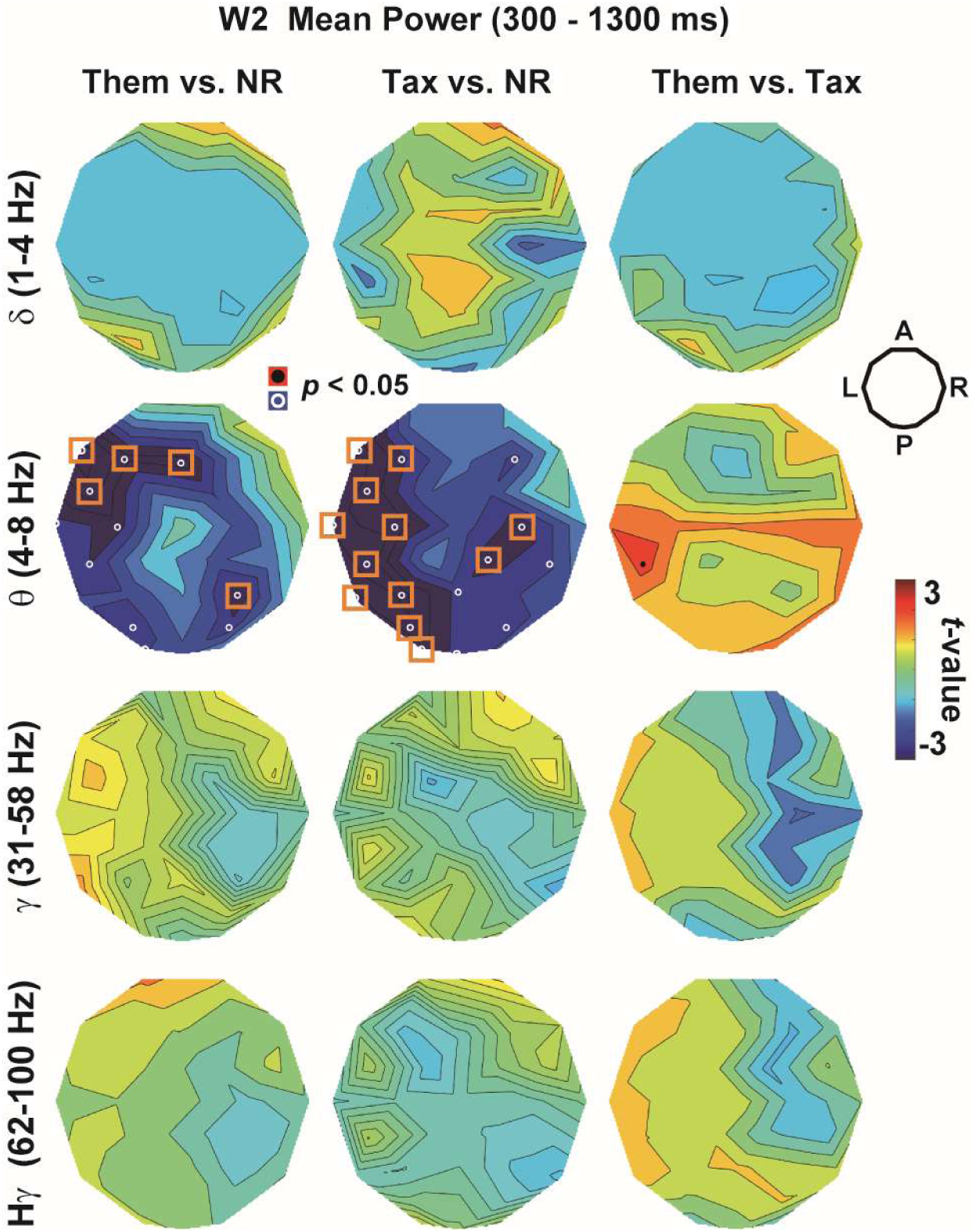
The *t*-maps of mean power in Experiment 2 in delta, theta, gamma, and high-gamma bands. No sensor showed significant difference in a direct comparison of Them vs. Tax (right panels).

**Figure S6.**
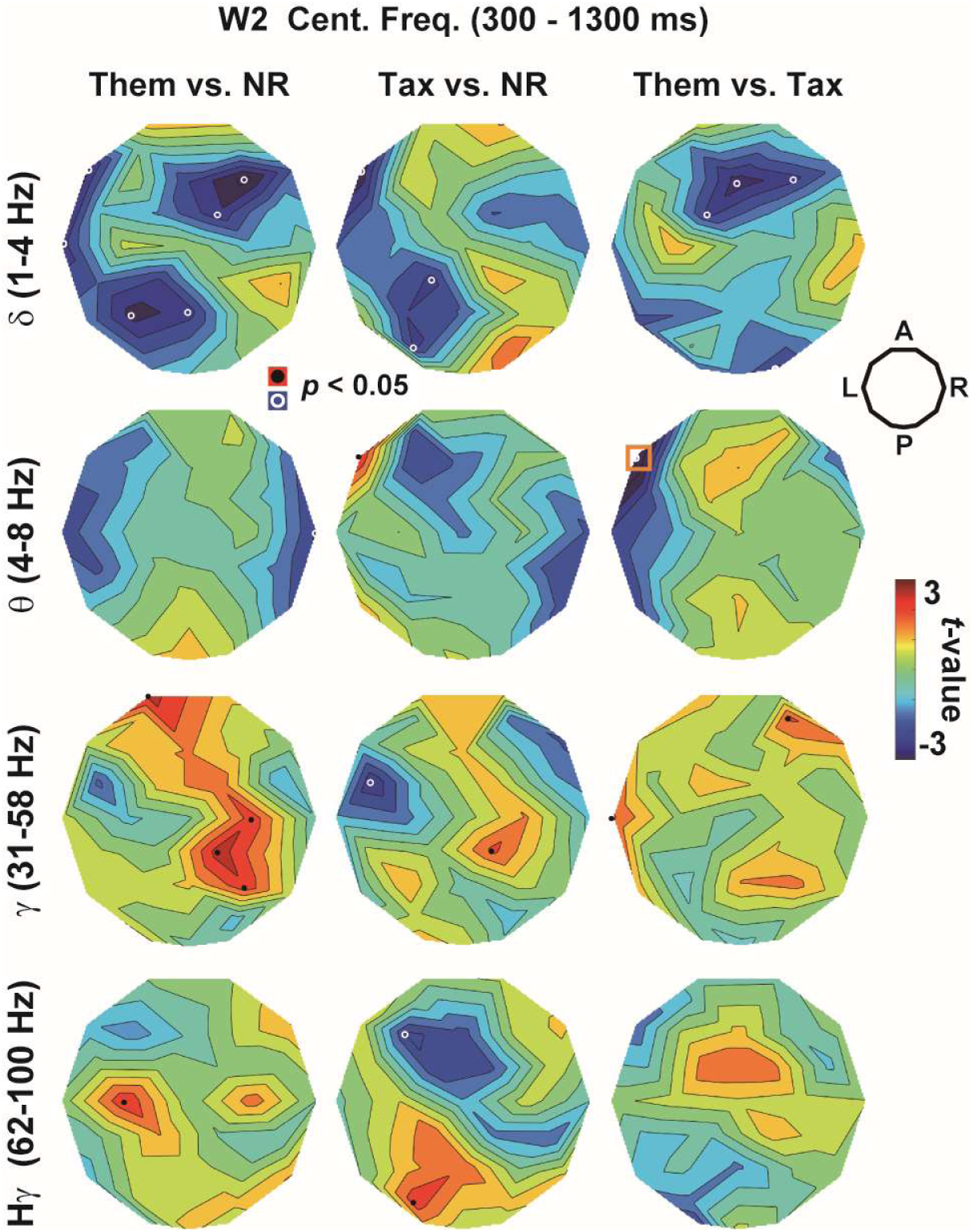
The *t*-maps of central frequency in Experiment 2 in delta, theta, gamma, and high-gamma bands. No sensor showed a significant difference after the correction of multiple comparisons, except for F7 (Them vs. Tax) in theta band.

## References

Adezati E, Liu X, Ding J, Thye M, Szaflarski JP, Mirman D (2024) Phase synchronization during the processing of taxonomic and thematic relations. Brain Lang 249:105379.

Anderson AJ, Binder JR, Fernandino L, Humphries CJ, Conant LL, Aguilar M, Wang X, Doko D, Raizada RDS (2017) Predicting Neural Activity Patterns Associated with Sentences Using a Neurobiologically Motivated Model of Semantic Representation. Cereb Cortex 27:4379–4395.

Benjamini Y, Hochberg Y (1995) Controlling the False Discovery Rate - a Practical and Powerful Approach to Multiple Testing. J Roy Stat Soc B Met 57:289–300.

Binder JR, Desai RH, Graves WW, Conant LL (2009) Where is the semantic system? A critical review and meta-analysis of 120 functional neuroimaging studies. Cereb Cortex 19:2767–2796.

Blackett DS, Varkey J, Wilmskoetter J, Roth R, Andrews K, Busby N, Gleichgerrcht E, Desai RH, Riccardi N, Basilakos A, Johnson LP, Kristinsson S, Johnson L, Rorden C, Spell LA, Fridriksson J, Bonilha L (2022) Neural network bases of thematic semantic processing in language production. Cortex 156:126–143.

Bondi E, Ding Y, Zhang Y, Maggioni E, He B (2025) Investigating the neurovascular coupling across multiple motor execution and imagery conditions: a whole-brain EEG-informed fMRI analysis. Neuroimage 317:121311.

Brainard DH (1997) The Psychophysics Toolbox. Spat Vis 10:433–436.

Budanitsky A, Hirst G (2006) Evaluating WordNet-based measures of lexical semantic relatedness. Comput Linguist 32:13–47.

Budel G, Jin Y, Van Mieghem P, Kitsak M (2023) Topological properties and organizing principles of semantic networks. Sci Rep 13:11728.

Carota F, Nili H, Pulvermuller F, Kriegeskorte N (2021) Distinct fronto-temporal substrates of distributional and taxonomic similarity among words: evidence from RSA of BOLD signals. Neuroimage 224:117408.

Chiang JN, Peng Y, Lu H, Holyoak KJ, Monti MM (2021) Distributed Code for Semantic Relations Predicts Neural Similarity during Analogical Reasoning. J Cogn Neurosci 33:377–389.

Chou TL, Wong CH, Chen SY, Fan LY, Booth JR (2019) Developmental changes of association strength and categorical relatedness on semantic processing in the brain. Brain Lang 189:10–19.

Cohen MX (2014) Fluctuations in oscillation frequency control spike timing and coordinate neural networks. Journal of Neuroscience 34:8988–8998.

Collins AM, Loftus EF (1975) Spreading Activation Theory of Semantic Processing. Psychol Rev 82:407–428.

Crutch SJ, Williams P, Ridgway GR, Borgenicht L (2012) The role of polarity in antonym and synonym conceptual knowledge: evidence from stroke aphasia and multidimensional ratings of abstract words. Neuropsychologia 50:2636–2644.

de Zubicaray GI, Hansen S, McMahon KL (2013) Differential processing of thematic and categorical conceptual relations in spoken word production. J Exp Psychol Gen 142:131–142.

Diedrich A, Arif Y, Taylor BK, Shen Z, Astorino PM, Lee WH, McCreery RW, Heinrichs-Graham E (2025) Distinct age-related alterations in alpha-beta neural oscillatory activity during verbal working memory encoding in children and adolescents. J Physiol 603:2387–2408.

Faul F, Erdfelder E, Lang AG, Buchner A (2007) G*Power 3: a flexible statistical power analysis program for the social, behavioral, and biomedical sciences. Behav Res Methods 39:175–191.

Fedorenko E, Ivanova AA, Regev TI (2024) The language network as a natural kind within the broader landscape of the human brain. Nat Rev Neurosci 25:289–312.

Fernandino L, Tong JQ, Conant LL, Humphries CJ, Binder JR (2022) Decoding the information structure underlying the neural representation of concepts. Proc Natl Acad Sci U S A 119.

Fu Z, Wang X, Wang X, Yang H, Wang J, Wei T, Liao X, Liu Z, Chen H, Bi Y (2023) Different computational relations in language are captured by distinct brain systems. Cereb Cortex 33:997–1013.

Goto H, Adachi T, Utsunomiya T, Chen IC (1979). Late positive component (LPC) and CNV during processing of linguistic information. In Human evoked potentials, D. Lehmann, and E. Callaway, eds. (New York, Plenum).

Hagoort P (2020) The meaning-making mechanism(s) behind the eyes and between the ears. Philos Trans R Soc Lond B Biol Sci 375:20190301.

Hall MC, Rempe MP, Casagrande CC, Glesinger RJ, Petro NM, Garrison GM, John JA, Dietz SM, Schantell M, Bai H, Arif Y, Embury CM, Bashford S, Okelberry HJ, Petts AJ, Keifer EL, May-Weeks PE, Picci G, Heinrichs-Graham E, Wilson TW (2025) Age-related alterations in alpha and beta oscillations support preservation of semantic processing in healthy aging. NPJ Aging 11:73.

Herlin B, Navarro V, Dupont S (2021) The temporal pole: From anatomy to function-A literature appraisal. J Chem Neuroanat 113:101925.

Hill F, Reichart R, Korhonen A (2015) SimLex-999: Evaluating Semantic Models With (Genuine) Similarity Estimation. Comput Linguist 41:665–695.

Holyoak KJ, Monti MM (2021) Relational Integration in the Human Brain: A Review and Synthesis. J Cogn Neurosci 33:341–356.

Hutchison KA (2003) Is semantic priming due to association strength or feature overlap?: A microanalytic review. Psychon Bull Rev 10:785–813.

Huth AG, de Heer WA, Griffiths TL, Theunissen FE, Gallant JL (2016) Natural speech reveals the semantic maps that tile human cerebral cortex. Nature 532:453–458.

Jackson RL, Hoffman P, Pobric G, Lambon Ralph MA (2015) The Nature and Neural Correlates of Semantic Association versus Conceptual Similarity. Cereb Cortex 25:4319–4333.

Jamali M, Grannan B, Cai J, Khanna AR, Munoz W, Caprara I, Paulk AC, Cash SS, Fedorenko E, Williams ZM (2024) Semantic encoding during language comprehension at single-cell resolution. Nature 631:610–616.

Jefferies E, Thompson H, Cornelissen P, Smallwood J (2020) The neurocognitive basis of knowledge about object identity and events: dissociations reflect opposing effects of semantic coherence and control. Philos Trans R Soc Lond B Biol Sci 375:20190300.

Jeon HA, Lee KM, Kim YB, Cho ZH (2009) Neural substrates of semantic relationships: common and distinct left-frontal activities for generation of synonyms vs. antonyms. Neuroimage 48:449–457.

Kalenine S, Peyrin C, Pichat C, Segebarth C, Bonthoux F, Baciu M (2009) The sensory-motor specificity of taxonomic and thematic conceptual relations: a behavioral and fMRI study. Neuroimage 44:1152–1162.

Kapur N, Ellison D, Smith MP, McLellan DL, Burrows EH (1992) Focal retrograde amnesia following bilateral temporal lobe pathology. A neuropsychological and magnetic resonance study. Brain 115 Pt 1:73–85.

Kliegr T, Zamazal O (2018) Antonyms are similar: Towards paradigmatic association approach to rating similarity in SimLex-999 and WordSim-353. Data & Knowledge Engineering 115:174–193.

Klimesch W, Schimke H, Pfurtscheller G (1993) Alpha frequency, cognitive load and memory performance. Brain Topogr 5:241–251.

Kuchinke L, Meer E, Krueger F (2009) Differences in processing of taxonomic and sequential relations in semantic memory: an fMRI investigation. Brain Cogn 69:245–251.

Kumar U (2018) The neural realm of taxonomic and thematic relation: an fMRI study. Language Cognition and Neuroscience 33:648–658.

Lau EF, Phillips C, Poeppel D (2008) A cortical network for semantics: (de)constructing the N400. Nat Rev Neurosci 9:920–933.

Lewis GA, Poeppel D, Murphy GL (2015) The neural bases of taxonomic and thematic conceptual relations: an MEG study. Neuropsychologia 68:176–189.

Li DG, Zhang XN, Wang GY (2011) Senior Chinese high school students’ awareness of thematic and taxonomic relations in L1 and L2. Bilingualism-Language and Cognition 14:444–457.

Li J, Pylkkanen L (2021) Disentangling Semantic Composition and Semantic Association in the Left Temporal Lobe. Journal of Neuroscience 41:6526–6538.

Li Y, Noguchi Y (2025) The role of beta band phase resetting in audio-visual temporal order judgment. Cogn Neurodyn 19:28.

Liu CY, Qin L, Tao R, Deng W, Jiang T, Wang N, Matthews S, Siok WT (2024) Delineating Region-Specific contributions and connectivity patterns for semantic association and categorization through ROI and Granger causality analysis. Brain Lang 258:105476.

Lu H, Chen D, Holyoak KJ (2012) Bayesian analogy with relational transformations. Psychol Rev 119:617–648.

Lucas M (2000) Semantic priming without association: A meta-analytic review. Psychonomic Bulletin & Review 7:618–630.

Meltzer JA, Kielar A, Panamsky L, Links KA, Deschamps T, Leigh RC (2017) Electrophysiological signatures of phonological and semantic maintenance in sentence repetition. Neuroimage 156:302–314.

Mirman D, Landrigan JF, Britt AE (2017) Taxonomic and thematic semantic systems. Psychol Bull 143:499–520.

Mummery CJ, Patterson K, Price CJ, Ashburner J, Frackowiak RS, Hodges JR (2000) A voxel-based morphometry study of semantic dementia: relationship between temporal lobe atrophy and semantic memory. Ann Neurol 47:36–45.

Murta T, Leite M, Carmichael DW, Figueiredo P, Lemieux L (2015) Electrophysiological correlates of the BOLD signal for EEG-informed fMRI. Hum Brain Mapp 36:391–414.

Nestor PJ, Fryer TD, Hodges JR (2006) Declarative memory impairments in Alzheimer’s disease and semantic dementia. Neuroimage 30:1010–1020.

Noguchi Y (2024) Harmonic memory signals in the human cerebral cortex induced by semantic relatedness of words. NPJ Sci Learn 9:6.

Noguchi Y, Inui K, Kakigi R (2004) Temporal dynamics of neural adaptation effect in the human visual ventral stream. Journal of Neuroscience 24:6283–6290.

Noguchi Y, Kakigi R (2020) Temporal codes of visual working memory in the human cerebral cortex: Brain rhythms associated with high memory capacity. Neuroimage 222:117294.

Oldfield RC (1971) The assessment and analysis of handedness: the Edinburgh inventory. Neuropsychologia 9:97–113.

Pelli DG (1997) The VideoToolbox software for visual psychophysics: transforming numbers into movies. Spat Vis 10:437–442.

Piantadosi ST, Muller DCY, Rule JS, Kaushik K, Gorenstein M, Leib ER, Sanford E (2024) Why concepts are (probably) vectors. Trends Cogn Sci 28:844–856.

Popov V, Hristova P, Anders R (2017) The relational luring effect: Retrieval of relational information during associative recognition. J Exp Psychol Gen 146:722–745.

Proskovec AL, Heinrichs-Graham E, Wilson TW (2019) Load modulates the alpha and beta oscillatory dynamics serving verbal working memory. Neuroimage 184:256–265.

Quillian MR (1967) Computers in Behavioral Science - Word Concepts - a Theory and Simulation of Some Basic Semantic Capabilities. Behavioral Science 12:410–430.

Ralph MA, Jefferies E, Patterson K, Rogers TT (2017) The neural and computational bases of semantic cognition. Nat Rev Neurosci 18:42–55.

Rempe MP, Casagrande CC, Embury CM, Arif Y, Bashford S, Garrison GM, Glesinger RJ, Okelberry HJ, Keifer EL, Picci G, Heinrichs-Graham E, Wilson TW (2025) Hippocampal and cortical oscillatory dynamics reflect semantic processing and predict behavioural performance. J Physiol 603:3089–3106.

Riccardi N, Schwen Blackett D, Broadhead A, den Ouden D, Rorden C, Fridriksson J, Bonilha L, Desai RH (2024) A Rose by Any Other Name: Mapping Taxonomic and Thematic Naming Errors Poststroke. J Cogn Neurosci 36:2251–2267.

Roehm D, Bornkessel-Schlesewsky I, Rosler F, Schlesewsky M (2007) To predict or not to predict: influences of task and strategy on the processing of semantic relations. J Cogn Neurosci 19:1259–1274.

Sachs O, Weis S, Zellagui N, Sass K, Huber W, Zvyagintsev M, Mathiak K, Kircher T (2011) How different types of conceptual relations modulate brain activation during semantic priming. J Cogn Neurosci 23:1263–1273.

Samenko I, Tikhonov A, Yamshchikov IP (2021) Intuitive Contrasting Map for Antonym Embeddings. arXiv 2004:12835.

Samuel IBH, Wang C, Hu ZH, Ding MZ (2018) The frequency of alpha oscillations: Task-dependent modulation and its functional significance. Neuroimage 183:897–906.

Scheible S, im Walde SS, Springorum S (2013) Uncovering Distributional Differences between Synonyms and Antonyms in aWord Space Model. Proceedings of the Sixth International Joint Conference on Natural Language Processing:489–497.

Schwartz MF, Kimberg DY, Walker GM, Brecher A, Faseyitan OK, Dell GS, Mirman D, Coslett HB (2011) Neuroanatomical dissociation for taxonomic and thematic knowledge in the human brain. Proc Natl Acad Sci U S A 108:8520–8524.

Setton R, Sheldon S, Turner GR, Spreng RN (2022) Temporal pole volume is associated with episodic autobiographical memory in healthy older adults. Hippocampus 32:373–385.

Snodgrass JG, Corwin J (1988) Pragmatics of measuring recognition memory: applications to dementia and amnesia. J Exp Psychol Gen 117:34–50.

Sowden S, Wright GR, Banissy MJ, Catmur C, Bird G (2015) Transcranial Current Stimulation of the Temporoparietal Junction Improves Lie Detection. Curr Biol 25:2447–2451.

Stochel JF, Sandberg CW (2025) Functional magnetic resonance imaging of taxonomic and thematic processing of abstract and concrete word pairs. J Neuropsychol 19:541–558.

Tadel F, Baillet S, Mosher JC, Pantazis D, Leahy RM (2011) Brainstorm: a user-friendly application for MEG/EEG analysis. Comput Intell Neurosci 2011:879716.

Teige C, Mollo G, Millman R, Savill N, Smallwood J, Cornelissen PL, Jefferies E (2018) Dynamic semantic cognition: Characterising coherent and controlled conceptual retrieval through time using magnetoencephalography and chronometric transcranial magnetic stimulation. Cortex 103:329–349.

Thatcher RW (1977). Evoked potential correlates of hemispheric lateralization during semantic information processing. In Lateralization in the nervous system, S. Harnad, R.W. Doty, L. Goldstein, J. Jaynes, and G. Krauthamer, eds. (New York, Academic Press).

Thye M, Geller J, Szaflarski JP, Mirman D (2021) Intracranial EEG evidence of functional specialization for taxonomic and thematic relations. Cortex 140:40–50.

Vaughan J, Sherif K, O’Sullivan RL, Herrmann DJ, Weldon DA (1982) Cortical evoked responses to synonyms and antonyms. Mem Cognit 10:225–231.

Vogel EK, Machizawa MG (2004) Neural activity predicts individual differences in visual working memory capacity. Nature 428:748–751.

Wang WC, Hsieh LT, Swamy G, Bunge SA (2021) Transient Neural Activation of Abstract Relations on an Incidental Analogy Task. J Cogn Neurosci 33:77–88.

Willners C (2001) Antonyms in Context : A Corpus-Based Semantic Analysis of Swedish Descriptive Adjectives. Travaux de l’Institut de Linguistique de Lund Lund University, Lund, Sweden 40.

Xavier M, Esteves I, Jorge J, Abreu R, Giraud AL, Sadaghiani S, Wirsich J, Figueiredo P (2025) Consistency of resting-state correlations between fMRI networks and EEG band power. Imaging Neuroscience 3.

Xu YW, Wang XS, Wang XY, Men WW, Gao JH, Bi YC (2018) Doctor, Teacher, and Stethoscope: Neural Representation of Different Types of Semantic Relations. Journal of Neuroscience 38:3303–3317.

Zhang Y, Mirman D, Hoffman P (2023) Taxonomic and thematic relations rely on different types of semantic features: Evidence from an fMRI meta-analysis and a semantic priming study. Brain Lang 242:105287.

